## Supplementary figures for "A phylogenetically-conserved axis of thalamocortical connectivity in the human brain"

Stuart Oldham[1,2] & Gareth Ball[1,3]

1. Developmental Imaging, Murdoch Children's Research Institute, The Royal Children's Hospital Melbourne, Australia
2. The Turner Institute for Brain and Mental Health, School of Psychological Sciences and Monash Biomedical Imaging, Monash University, Clayton, Australia
3. Department of Paediatrics, University of Melbourne, Melbourne, Australia

**Corresponding author**
Dr Stuart Oldham
Developmental Imaging,
Murdoch Children's Research Institute
Parkville 3052 VIC
Australia


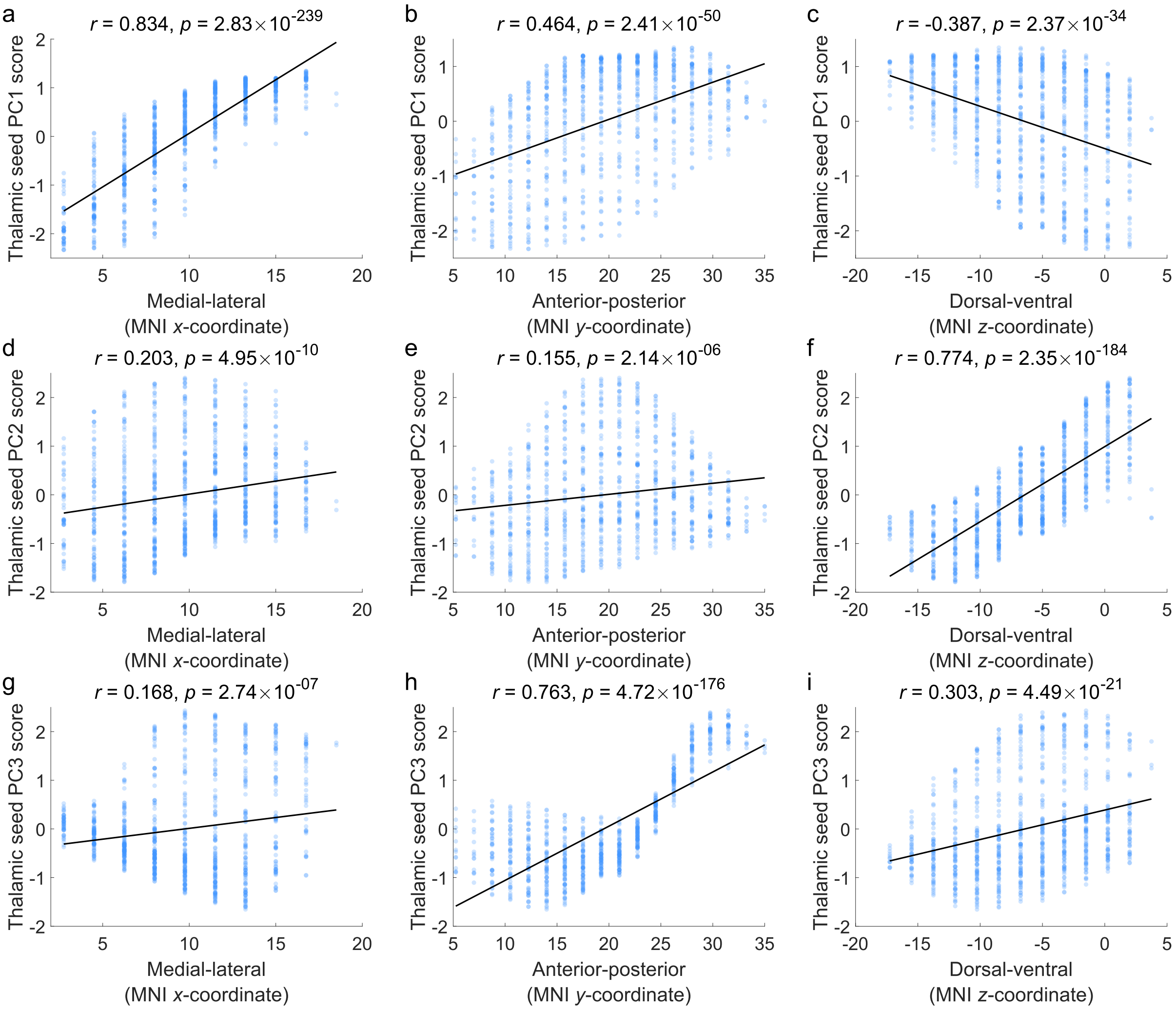


**Fig. S1. Correlations between seed PC1-PC3 score and cartesian axes position. a**, PC1 score with medial-lateral axis (MNI *x*-coordinate; Pearson's $r(919) = 0.834$, $p = 2.83 \times 10^{-239}$, CI = $[0.813, 0.852]$, two-tailed). **b**, PC1 score with anterior-posterior axis (MNI *y*-coordinate; Pearson's $r(919) = 0.464$, $p = 2.41 \times 10^{-45}$, CI = $[0.412, 0.513]$, two-tailed). **c**, PC1 score with dorsal-ventral axis (MNI *z*-coordinate; Pearson's $r(919) = -0.387$, $p = 2.37 \times 10^{-34}$, CI = $[-0.441, -0.331]$, two-tailed). **d**, PC2 score with medial-lateral axis (MNI *x*-coordinate; Pearson's $r(919) = 0.203$, $p = 4.95 \times 10^{-10}$, CI = $[0.140, 0.264]$, two-tailed). **e**, PC2 score with anterior-posterior axis (MNI *y*-coordinate; Pearson's $r(919) = 0.155$, $p = 2.14 \times 10^{-06}$, CI = $[0.092, 0.218]$, two-tailed). **f**, PC2 score with dorsal-ventral axis (MNI *z*-coordinate; Pearson's $r(919) = 0.774$, $p = 2.35 \times 10^{-184}$, CI = $[0.746, 0.798]$, two-tailed. **g**, PC3 score with medial-lateral axis (MNI *x*-coordinate; Pearson's $r(919) = 0.169$, $p = 2.74 \times 10^{-07}$, CI = $[0.105, 0.231]$, two-tailed). **h**, PC3 score with anterior-posterior axis (MNI *y*-coordinate; Pearson's $r(919) = 0.763$, $p = 4.72 \times 10^{-176}$, CI = $[0.734, 0.788]$, two-tailed. **i**, PC3 score with dorsal-ventral axis (MNI *z*-coordinate; Pearson's $r(919) = 0.303$, $p = 4.49 \times 10^{-21}$, CI = $[0.244, 0.361]$, two-tailed). Note that the MNI coordinates have been adjusted so that they align with the orientation of a specific plane, with the values increasing in the direction of that plane's cardinal orientation (i.e., medial to lateral etc.). The black line indicates the line of best fit. Note that despite some relationships being non-linear, there was little difference in the correlation coefficient when a Spearman's correlation was used. Source data are provided as a Source Data file.

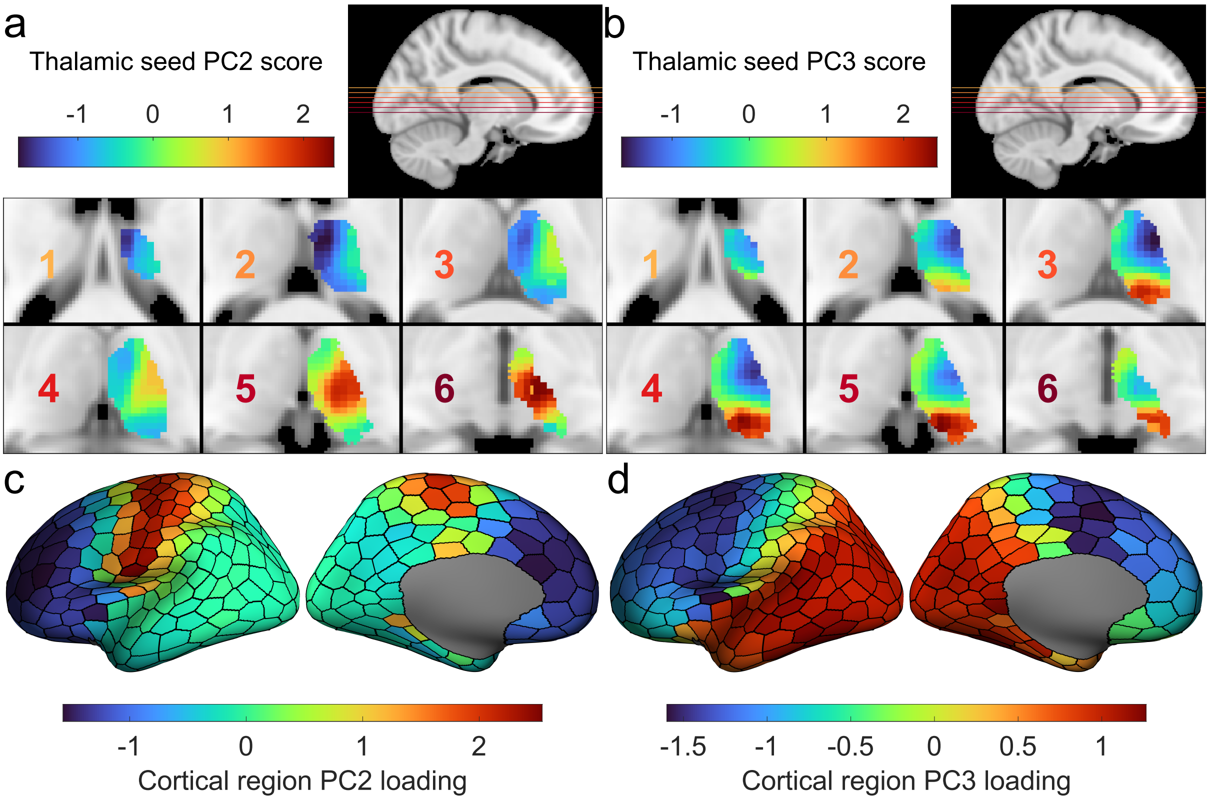


**Fig. S2. Spatial projections of PC2 and PC3. a**, Projection of PC2 scores of the human data onto thalamic voxels. PC2 scores for each seed are projected onto the closest voxels in the thalamic mask. **b**, Projection of PC3 scores of the human data onto thalamic voxels. PC2 scores for each seed are projected onto the closest voxels in the thalamic mask. **c**, PC2 loadings for cortical regions projected onto the cortical surface. **d**, PC3 loadings for cortical regions projected onto the cortical surface. Source data are provided as a Source Data file.

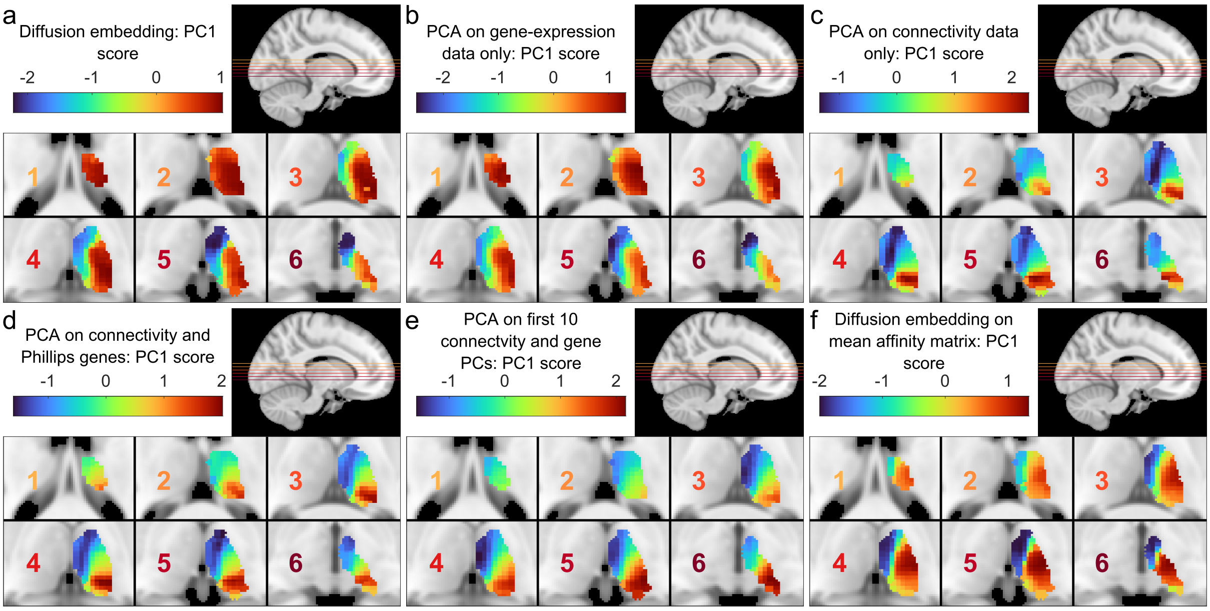


**Fig. S3. Alternative decompositions of thalamic gene expression and connectivity data.** The resulting PC1 scores projected onto the thalamus after **a**, Nonlinear diffusion embedding of combined gene expression and connectivity data. **b**, PCA on gene expression data only. **c**, PCA on connectivity only. **d**, PCA on expression data of genes identified in an independent study [Phillips et al.] and connectivity data. **e**, PCA on the first combined 10 components from separate PCAs on the gene expression and connectivity data **f**, Diffusion embedding applied to the average affinity matrices (determined by cosine angle) calculated separately for gene expression and connectivity data. Source data are provided as a Source Data file.

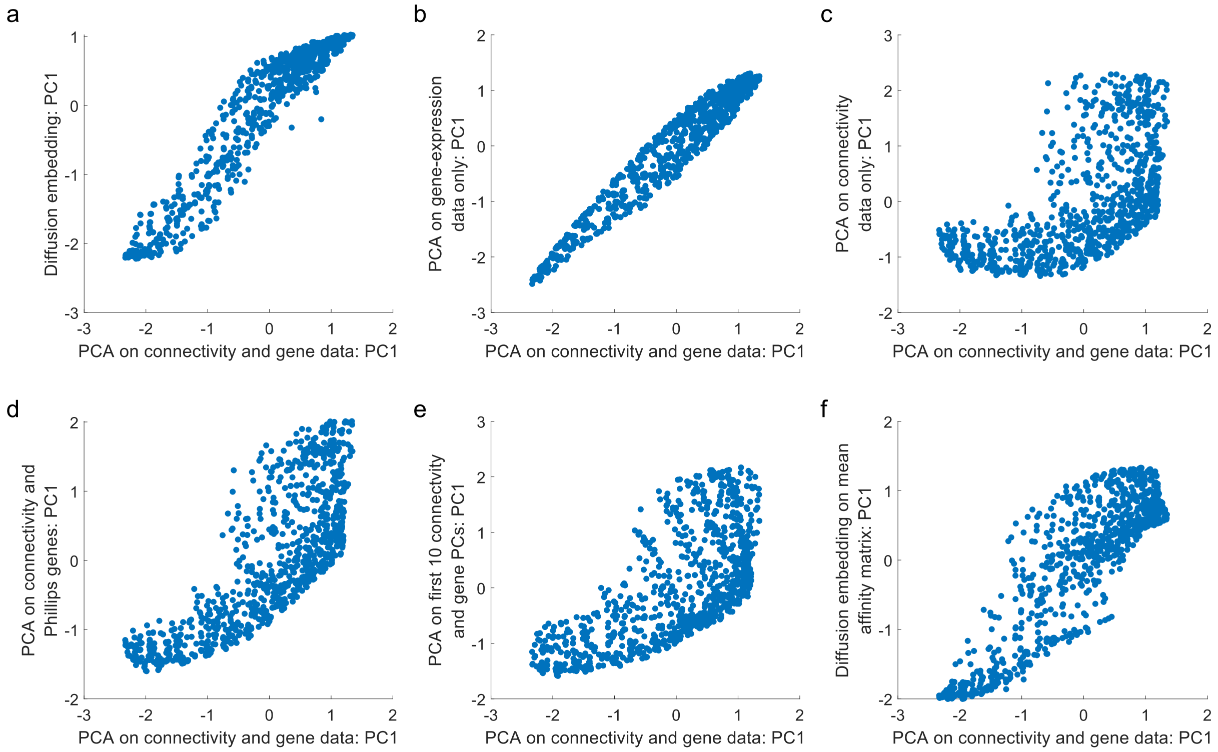


**Fig. S4. Comparison between original and alternative decompositions of thalamic gene expression and connectivity data.** Comparison between PC1 scores for the original (PCA on connectivity and gene data) and alternative decompositions **a**, Nonlinear diffusion embedding of combined gene expression and connectivity data. **b**, PCA on gene expression data only. **c**, PCA on connectivity only. **d**, PCA on expression data of genes identified in an independent study [Phillips et al.] and connectivity data. **e**, PCA on the first combined 10 components from separate PCAs on the gene expression and connectivity data **f**, Diffusion embedding applied to the average affinity matrices (determined by cosine angle) calculated separately for gene expression and connectivity data. Source data are provided as a Source Data file.

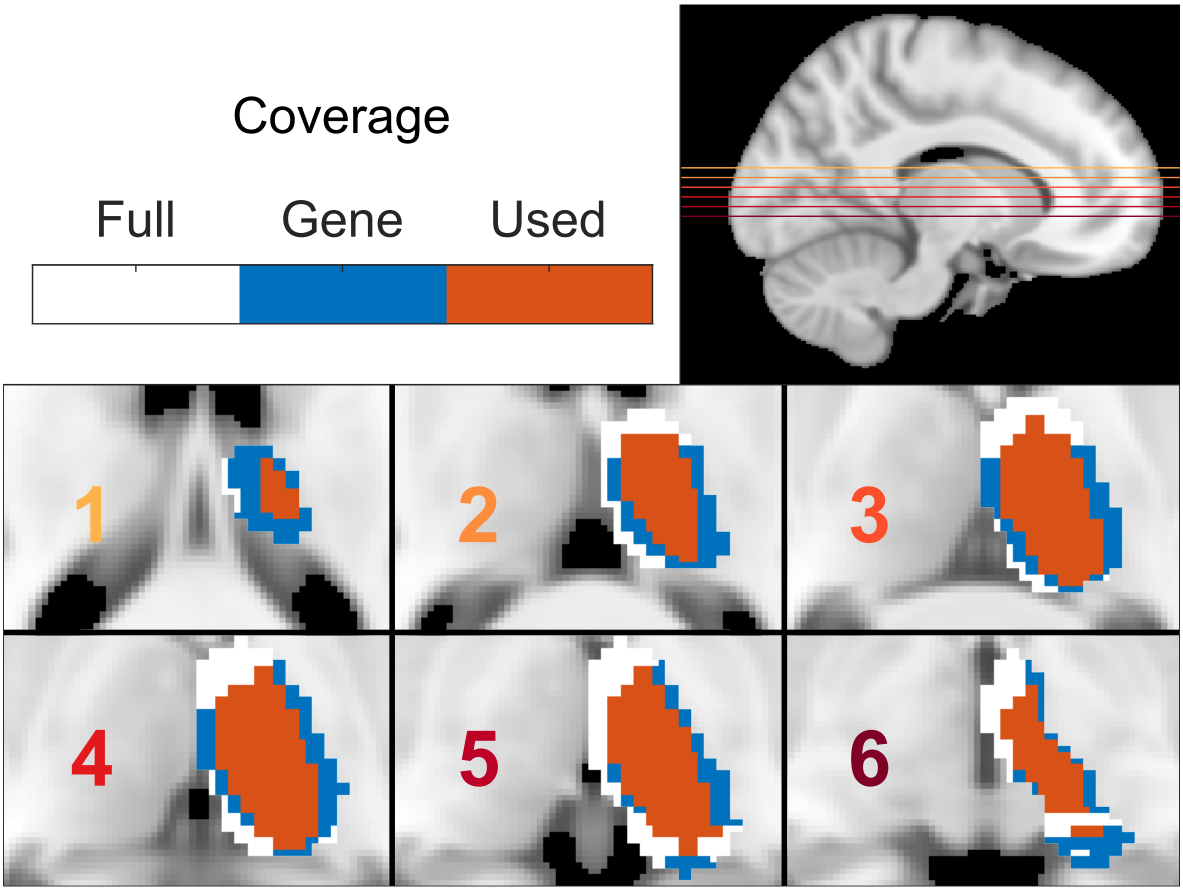


**Fig. S5. Coverage of seed points in the thalamus.** The full thalamic mask (white) is specified based on an atlas (Melbourne Subcortex Atlas). Within this mask, areas with gene expression are colored blue, whilst areas which were used in the main decomposition (i.e., areas which showed consistent registrations across individuals and had gene expression data available) are shown in orange. Anterior-medial areas of the thalamus lacked coverage. Source data are provided as a Source Data file.

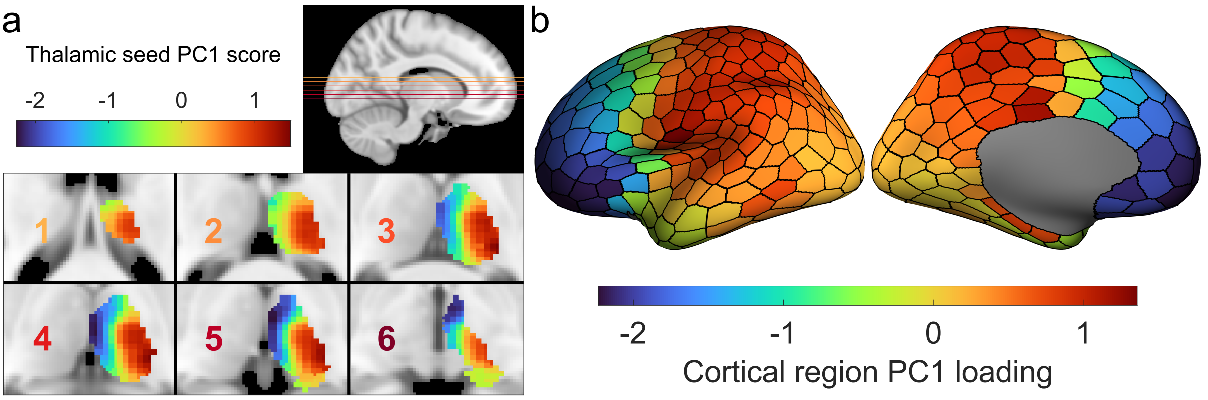


**Fig. S6. Decomposition results for PC1 when using all seeds with gene-expression information. a**, Projection of PC1 scores of the human data onto thalamic voxels. Scores for each seed are projected onto the closest voxels in the thalamic mask. **b**, PC1 loadings for cortical regions projected onto the cortical surface. Source data are provided as a Source Data file.

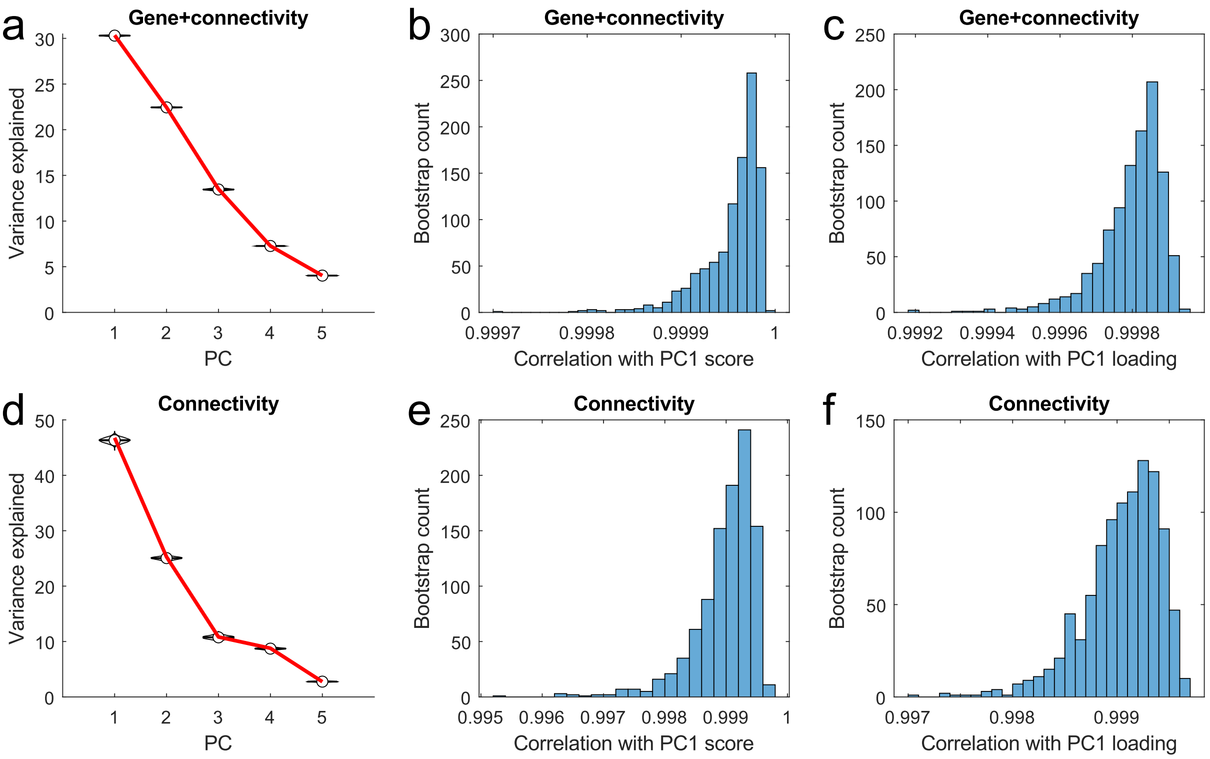


**Fig. S7. Bootstrapping results for the decompositions performed.** Bootstrapping (1000 iterations) was performed to obtain a new sampling of the connectivity/tractography data. This was either combined with the gene data (gene+connectivity) or used by itself (connectivity) and a PCA was performed and the result was compared to the respective original decomposition. **a**, Variance explained across the first five components for the decomposition of the gene and connectivity data. The red line indicates the original result, whilst the violin plots indicate variation in the variance explained across bootstrap iterations. **b**, Histogram of correlation values between the PC1 scores of the original decomposition and those obtained from each bootstrap iteration. **c**, Histogram of correlation values between the PC1 loadings of the original decomposition and those obtained from each bootstrap iteration. **d**, Variance explained across the first five components for the decomposition of the connectivity data. The red line indicates the original result, whilst the violin plots indicate variation in the variance explained across bootstrap iterations. **e**, Histogram of correlation values between the PC1 scores of the decomposition of the connectivity data and those obtained from each bootstrap iteration. **f**, Histogram of correlation values between the PC1 loadings of the decomposition of the connectivity data and those obtained from each bootstrap iteration. Source data are provided as a Source Data file.

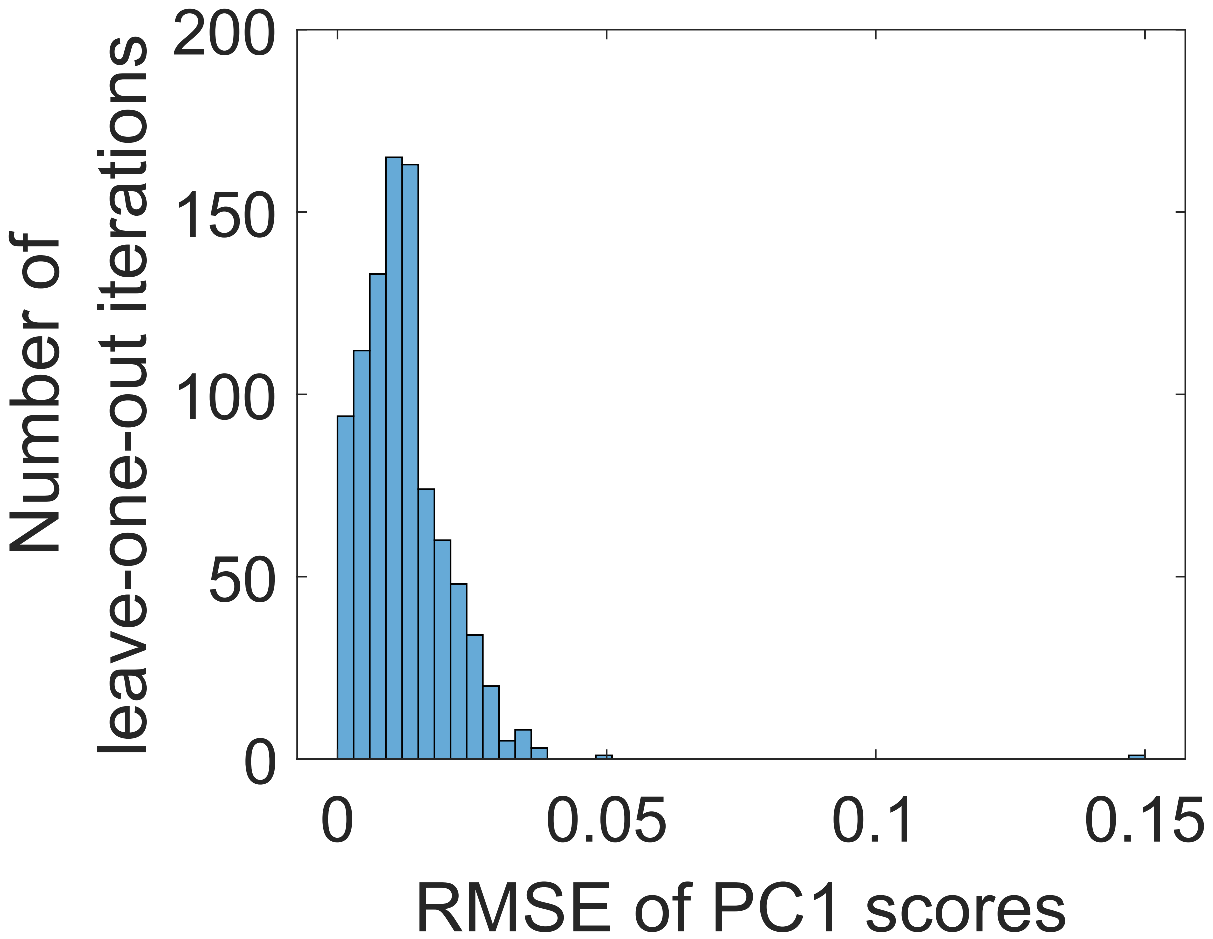


**Fig. S8. Leave-one-out cross-validation.** Leave-one-out cross-validation was performed by iteratively removing the data for a single seed from the concatenated connectivity and gene expression matrix. The decomposition was performed on this and the root-mean-square-error (RMSE) for the PC1 scores with the corresponding scores from the original decomposition was taken. The histogram shows the distribution of RMSE values for each iteration. Source data are provided as a Source Data file.

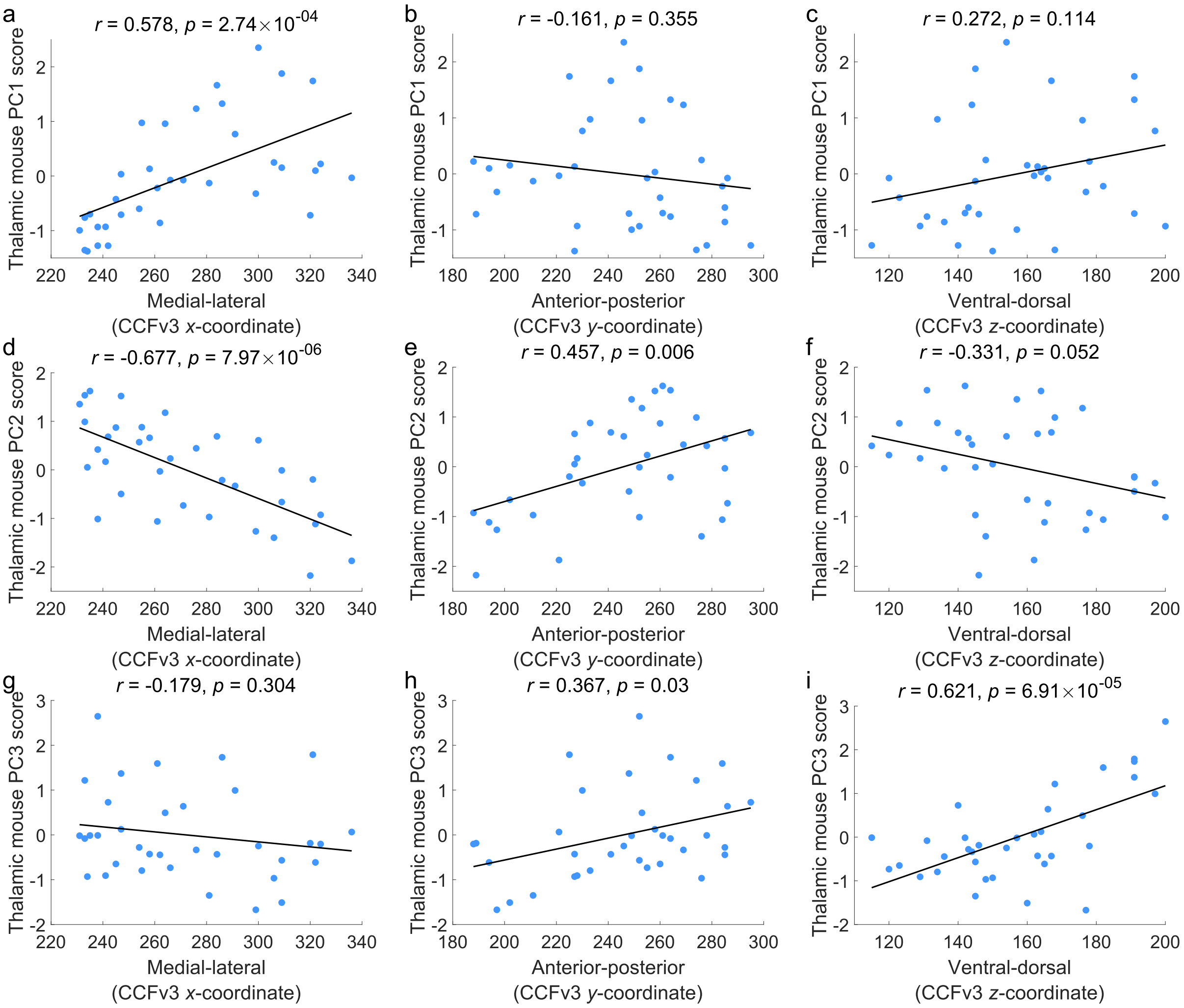


**Fig. S9. Correlations between thalamic mouse PC1-PC3 score and cartesian axes position. a**, mPC1 score with medial-lateral axis (Allen Mouse Brain Common Coordinate Framework version 3 [CCFv3] *x*-axis coordinate; Pearson's $r(33) = 0.578$, $p = 2.74 \times 10^{-04}$, CI = $[0.304, 0.764]$, two-tailed). **b**, mPC1 score with anterior-posterior axis (CCFv3 *y*-axis coordinate; Pearson's $r(33) = -0.161$, $p = 0.355$, CI = $[-0.469, 0.182]$, two-tailed). **c**, mPC1 score with dorsal-ventral axis (CCFv3 *z*-axis coordinate; Pearson's $r(34) = 0.272$, $p = 0.114$, CI = $[-0.068, 0.555]$, two-tailed). **d**, mPC2 score with medial-lateral axis (CCFv3 *x*-axis coordinate; Pearson's $r(33) = -0.677$, $p = 7.97 \times 10^{-06}$, CI = $[-0.824, -0.444]$, two-tailed). **e**, mPC2 score with anterior-posterior axis (CCFv3 *y*-axis coordinate; Pearson's $r(33) = 0.457$, $p = 0.006$, CI = $[0.146, 0.686]$, two-tailed). **f**, mPC2 score with dorsal-ventral axis (CCFv3 *z*-axis coordinate; Pearson's $r(33) = -0.331$, $p = 0.0518$, CI = $[-0.599, 0.002]$, two-tailed). **g**, mPC3 score with medial-lateral axis (CCFv3 *x*-axis coordinate; Pearson's $r(33) = -0.179$, $p = 0.304$, CI = $[-0.483, 0.164]$, two-tailed). **h**, mPC3 score with anterior-posterior axis (CCFv3 *y*-axis coordinate; Pearson's $r(33) = 0.367$, $p = 0.030$, CI = $[0.038, 0.624]$, two-tailed). **i**, mPC3 score with dorsal-ventral axis (CCFv3 *z*-axis coordinate; Pearson's $r(33) = 0.62087$, $p = 6.91 \times 10^{-05}$, CI = $[0.363, 0.791]$, two-tailed). The black line indicates the line of best fit. Source data are provided as a Source Data file.

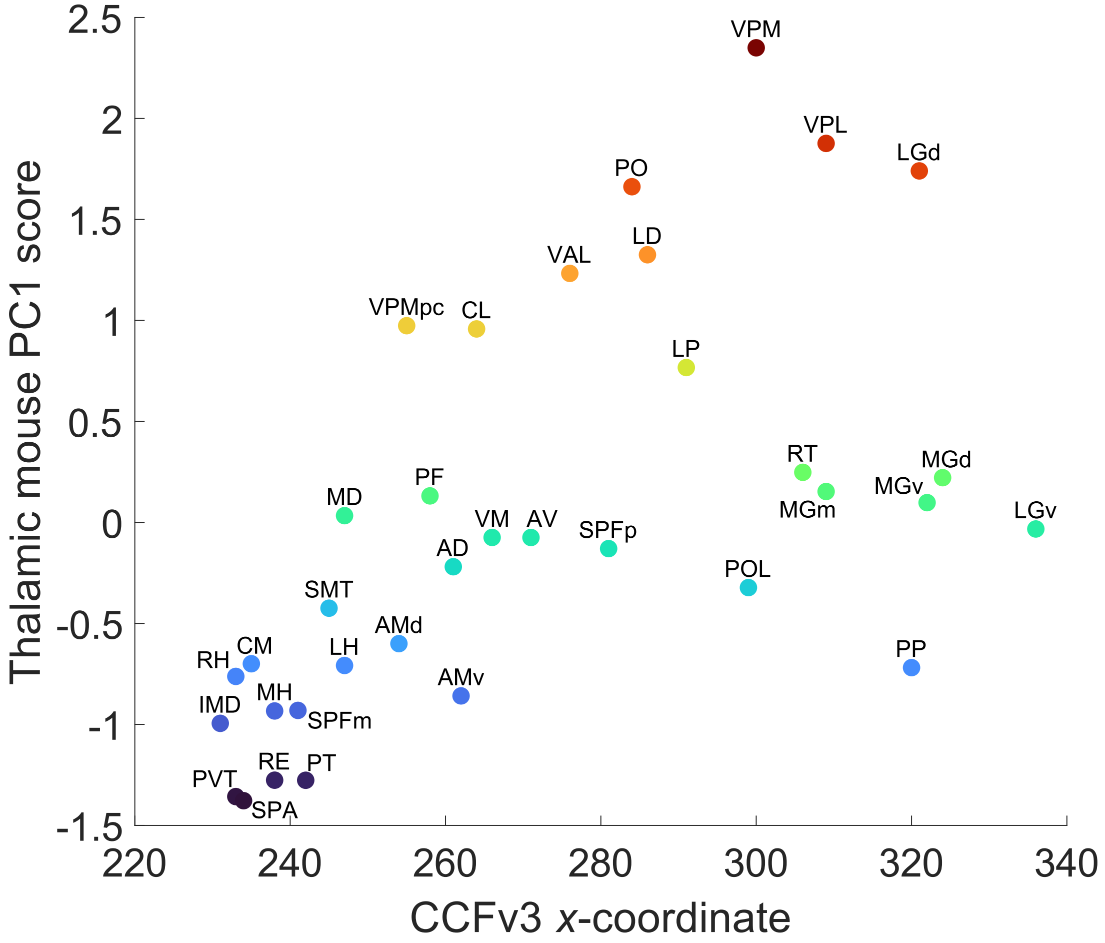


**Fig. S10. Relationship between thalamic nuclei PC1 score and medial-lateral position in the mouse**. The PC1 score for nuclei from the in the Allen Mouse Brain data (mPC1) is shown against its x-coordinate (representing medial-lateral position) in Allen Mouse Brain Common Coordinate Framework version 3 (CCFv3) space. Points are colored according to their PC1 score. AD = Anterodorsal nucleus, AMd = Anteromedial nucleus, dorsal part, AMv = Anteromedial nucleus, ventral part, AV = Anteroventral nucleus of thalamus, CL = Central lateral nucleus of the thalamus, CM = Central medial nucleus of the thalamus, IMD = Intermediodorsal nucleus of the thalamus, LD = Lateral dorsal nucleus of thalamus, LGd = Dorsal part of the lateral geniculate complex, LGv = Ventral part of the lateral geniculate complex, LH = Lateral habenula, LP = Lateral posterior nucleus of the thalamus, MD = Mediodorsal nucleus of thalamus, MGd = Medial geniculate complex, dorsal part, MGm = Medial geniculate complex, medial part, MGv = Medial geniculate complex, ventral part, MH = Medial habenula, PF = Parafascicular nucleus, PO = Posterior complex of the thalamus, POL = Posterior limiting nucleus of the thalamus, PP = Peripeduncular nucleus, PT = Parataenial nucleus, PVT = Paraventricular nucleus of the thalamus, RE = Nucleus of reuniens, RH = Rhomboid nucleus, RT = Reticular nucleus of the thalamus, SMT = Submedial nucleus of the thalamus, SPA = Subparafascicular area, SPFm = Subparafascicular nucleus, magnocellular part, SPFp = Subparafascicular nucleus, parvicellular part, VAL = Ventral anterior-lateral complex of the thalamus, VM = Ventral medial nucleus of the thalamus, VPL = Ventral posterolateral nucleus of the thalamus, VPM = Ventral posteromedial nucleus of the thalamus, VPMpc = Ventral posteromedial nucleus of the thalamus, parvicellular part. Source data are provided as a Source Data file.

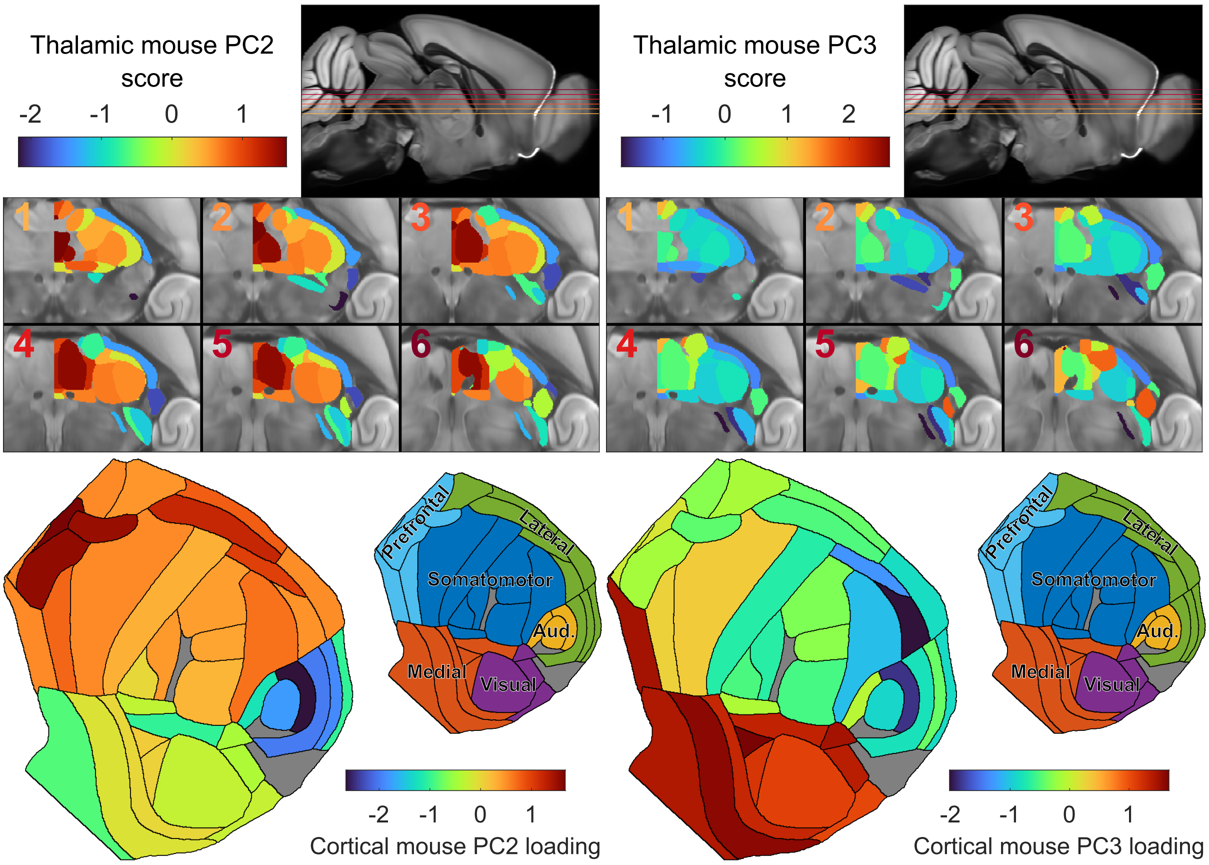


**Fig. S11. Spatial projections of mouse PC2 and PC3. a**, Projection of PC2 scores of the Allen Mouse Brain Atlas (AMBA) data (mPC2) onto respective thalamic nuclei. Note that while axonal-tracing data was obtained in the right hemisphere, we project the PC scores onto the left to enable straightforward comparison with the human data. **b**, Projection of PC3 scores of the AMBA data (mPC3) onto respective thalamic nuclei. **c**, Projection of PC2 loadings of the AMBA data (mPC2) onto respective cortical regions, displayed as a flat map. The smaller flat map plot indicates major cortical divisions (prefrontal, lateral, somatomotor, visual, medial, and auditory). Grey regions indicate cortical areas which no gene expression and/or connectivity data was available **d**, Projection of PC3 loadings of the AMBA data (mPC3) onto respective cortical regions, displayed as a flat map. Source data are provided as a Source Data file.

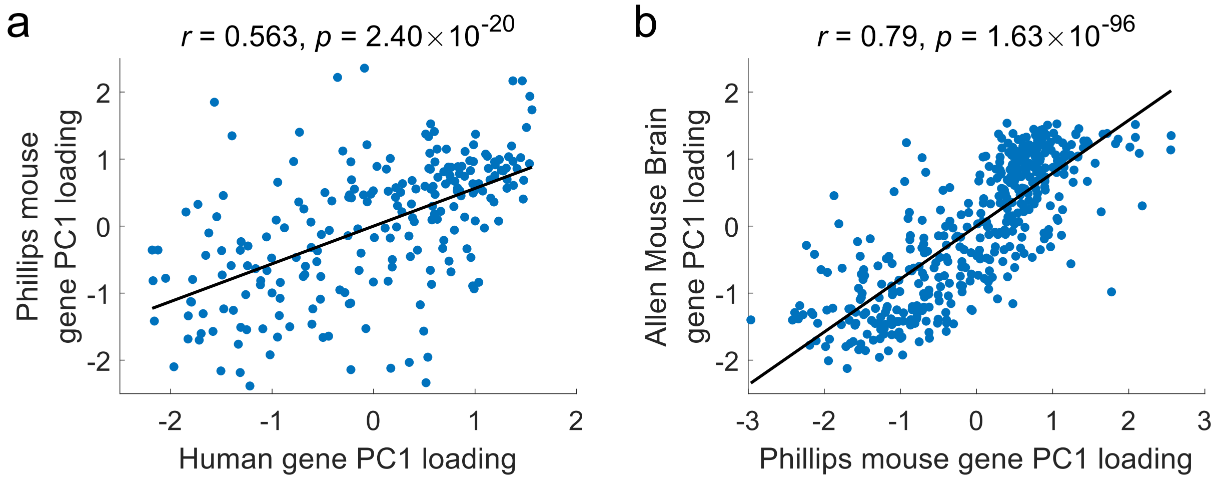


**Fig. S12. Relationships with Phillips et al. gene loadings. a**, Relationship between PC1 loadings for homolog genes in the human and Phillips et al. data (Pearson's $r(225) = 0.563$, $p = 2.40 \times 10^{-20}$, CI $= [0.467, 0.645]$, two-tailed). **b**, Relationship between PC1 loadings for homolog genes in the Allen Mouse Brain and Phillips et al. data (Pearson's $r(445) = 0.790$, $p = 1.63 \times 10^{-96}$, CI $= [0.752, 0.822]$, two-tailed). The black line indicates the line of best fit. Source data are provided as a Source Data file.

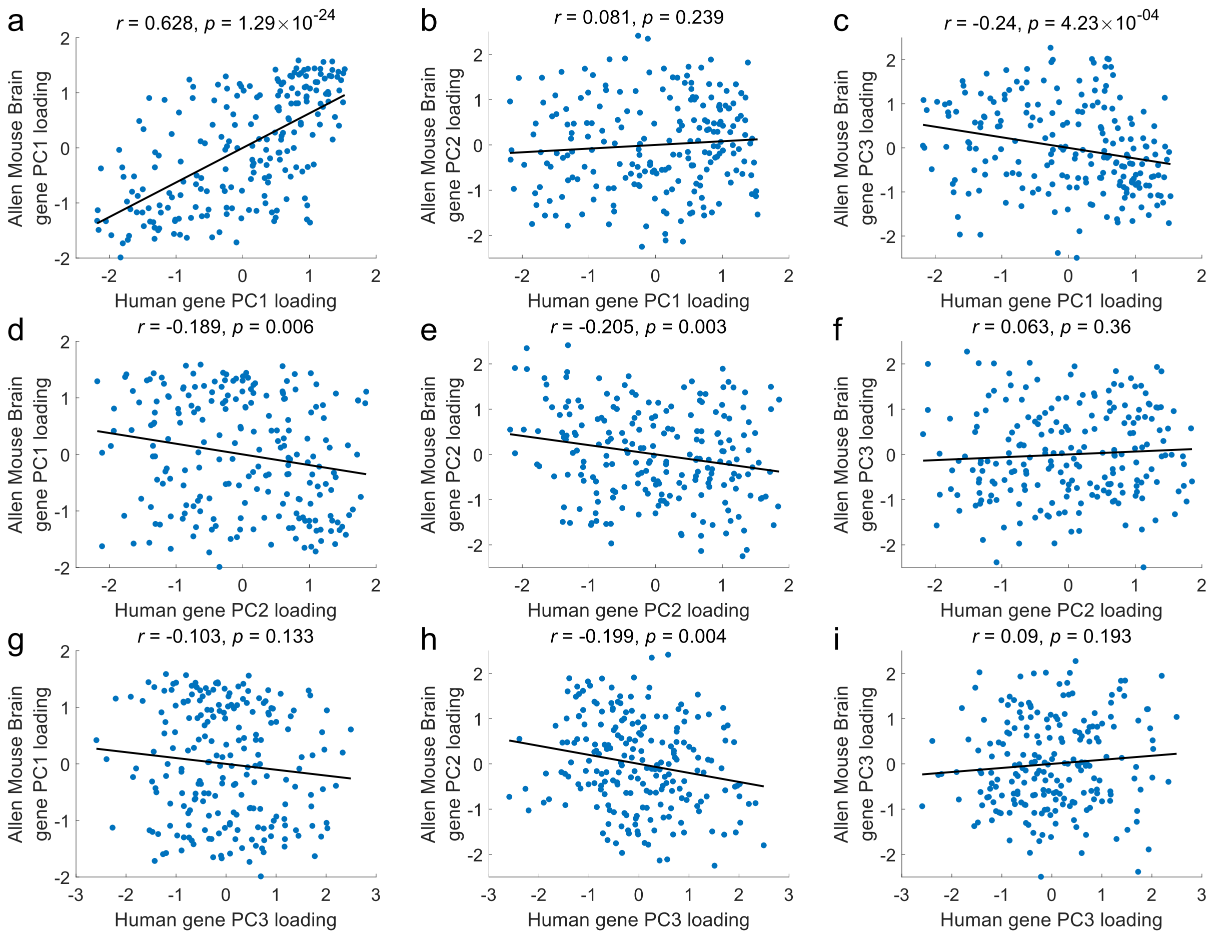


**Fig. S13. Relationships between mouse and human PC gene loadings. a**, Relationship between human PC1 and mouse PC1 loadings for homolog genes (Pearson's $r(210) = 0.628, p = 1.29 \times 10^{-24}$, CI = $[0.538, 0.703]$, two-tailed). **b**, Relationship between human PC1 and mouse PC2 loadings for homolog genes (Pearson's $r(210) = 0.081$, $p = 0.239$, CI = $[-0.054, 0.214]$, two-tailed). **c**, Relationship between human PC1 and mouse PC3 loadings for homolog genes (Pearson's $r(210) = -0.240$, $p = 0.0004$, CI = $[-0.363, -0.109]$, two-tailed). **d**, Relationship between human PC2 and mouse PC1 loadings for homolog genes (Pearson's $r(210) = -0.189$, $p = 0.006$, CI = $[-0.316, -0.056]$, two-tailed). **e**, Relationship between human PC2 and mouse PC2 loadings for homolog genes (Pearson's $r(210) = -0.205$, $p = 0.003$, CI = $[-0.330, -0.072]$, two-tailed). **f**, Relationship between human PC2 and mouse PC3 loadings for homolog genes (Pearson's $r(210) = 0.063, p = 0.360$, CI = $[-0.072, 0.196]$, two-tailed). **g**, Relationship between human PC3 and mouse PC1 loadings for homolog genes (Pearson's $r(210) = -0.103$, $p = 0.133$, CI = $[-0.235, 0.0318]$, two-tailed). **h**, Relationship between human PC3 and mouse PC2 loadings for homolog genes (Pearson's $r(210) = -0.199$, $p = 0.004$, CI = $[-0.325, -0.066]$, two-tailed). **i**, Relationship between human PC3 and mouse PC3 loadings for homolog genes (Pearson's $r(210) = 0.090$, $p = 0.193$, CI = $[-0.046, 0.222]$, two-tailed). The black line indicates the line of best fit. Source data are provided as a Source Data file.

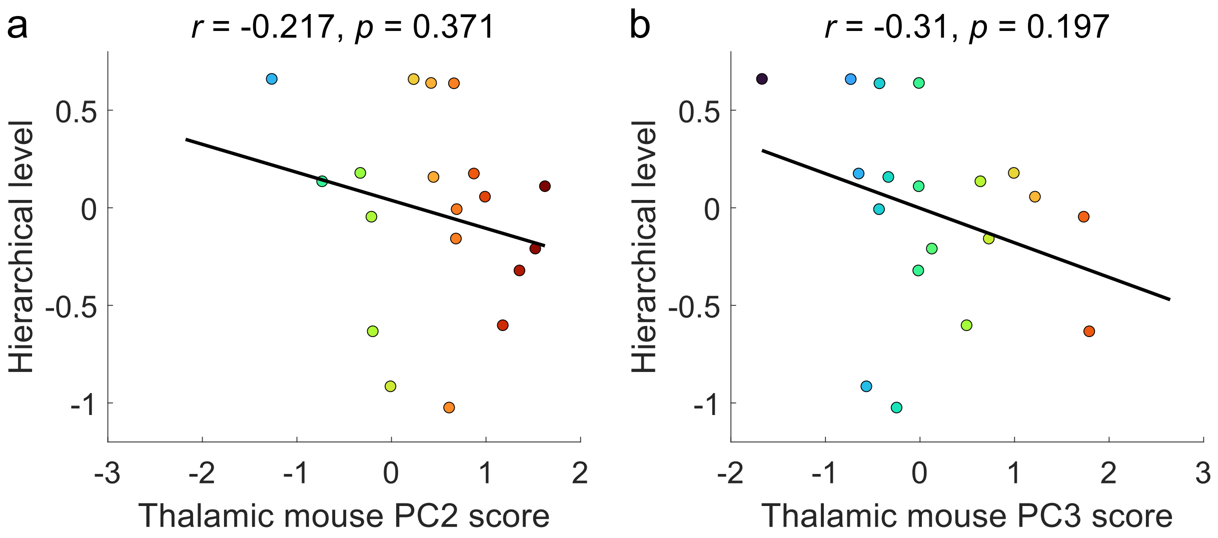


**Fig. S14. Relationship between mouse thalamic nuclei PC2 and PC3 scores and a measure of hierarchical organisation. a**, Relationship with thalamic mouse PC2 (mPC2) score (Pearson's $r(17) = -0.217$, $p = 0.371$, CI $= [-0.611, 0.263]$, two-tailed). **b**, Relationship with thalamic mouse PC3 (mPC3) score (Pearson's $r(17) = -0.310$, $p = 0.197$, CI $= [-0.670, 0.168]$, two-tailed). The black line indicates the line of best fit. Source data are provided as a Source Data file.

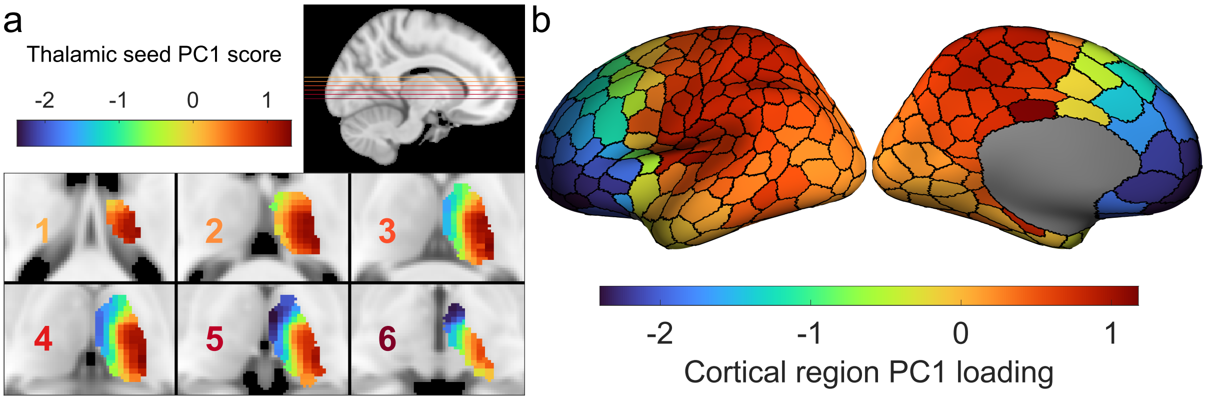


**Fig. S15. Decomposition results for PC1 when using the Schaefer 400 parcellation. a**, Projection of PC1 scores of the human data onto thalamic voxels. Scores for each seed are projected onto the closest voxels in the thalamic mask. **b**, PC1 loadings for cortical regions projected onto the cortical surface. Source data are provided as a Source Data file.

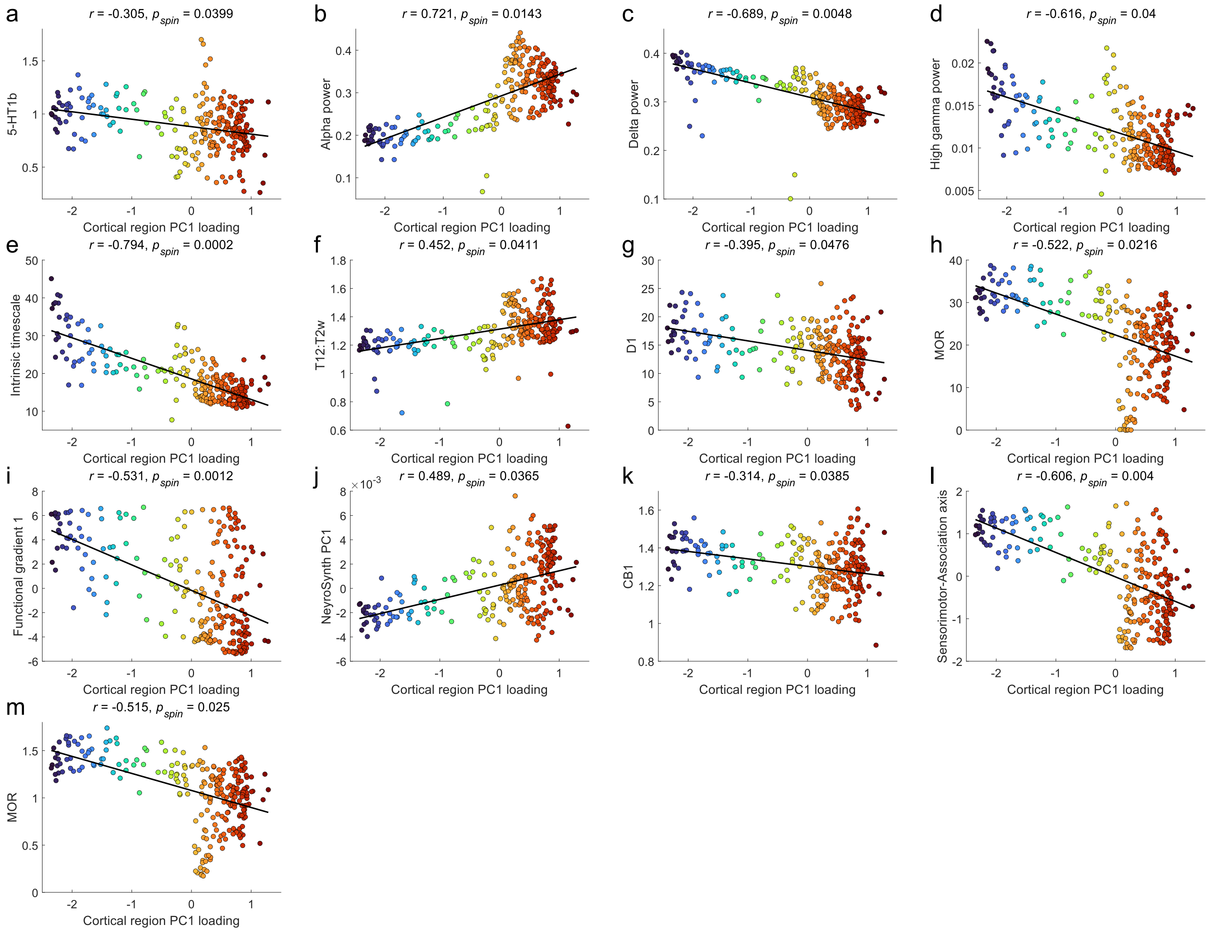


**Fig. S16. Significant relationships between PC1 cortical loadings and cortical measures from the *neuromaps* toolbox.** Points are coloured according to their cortical region PC1 loading. The black line indicates the line of best fit. Significant relationships were detected via a spin-test ($p_{spin} < 0.05$; two-tailed) for the following cortical features: **a**, PET tracer binding (BPnd) to 5-HT1b (serotonin receptor) (Pearson's $r(248) = -0.305$, $p_{spin} = 0.0399$, CI = $[-0.414, -0.188]$, two-tailed). **b**, MEG alpha (8-12 Hz) power distribution from the Human Connectome Project S1200 release (Pearson's $r(248) = 0.721$, $p_{spin} = 0.0143$, CI = $[0.655, 0.775]$, two-tailed). **c**, MEG delta (2-4 Hz) power distribution from the Human Connectome Project S1200 release (Pearson's $r(248) = -0.689$, $p_{spin} = 0.0048$, CI = $[-0.749, -0.618]$, two-tailed). **d**, MEG high gamma (60-90 Hz) power distribution from the Human Connectome Project S1200 release (Pearson's $r(248) = -0.616$, $p_{spin} = 0.04$, CI = $[-0.687, -0.532]$, two-tailed). **e**, MEG intrinsic timescale from the Human Connectome Project S1200 release (Pearson's $r(248) = -0.794$, $p_{spin} = 0.0002$, CI = $[-0.835, -0.743]$, two-tailed). **f**, MRI T1w/T2w ratio from the Human Connectome Project S1200 release (Pearson's $r(248) = 0.452$, $p_{spin} = 0.0411$, CI = $[0.347, 0.545]$, two-tailed). **g**, PET tracer binding (BPnd) to D1 (dopamine receptor; Pearson's $r(248) = -0.395$, $p_{spin} = 0.0476$, CI = $[-0.495, -0.285]$, two-tailed). **h**, PET tracer binding (BPnd) to MOR (mu-opioid receptor; Pearson's $r(248) = -0.522$, $p_{spin} = 0.0216$, CI = $[-0.607, -0.425]$, two-tailed). **i**, Diffusion map embedding gradient 1 of group-averaged functional connectivity (Pearson's $r(248) = -0.531$, $p_{spin} = 0.0012$, CI = $[-0.615, -0.436]$, two-tailed). **j**, PC1 of Neurosynth terms in the Cognitive Atlas (123 terms total; Pearson's $r(248) = 0.489$, $p_{spin} = 0.0365$, CI = $[0.388, 0.578]$, two-tailed). **k**, PET tracer binding (Vt) to CB1 (cannabinoid receptor; Pearson's $r(248) = -0.314$, $p_{spin} = 0.0385$, CI = $[-0.421, -0.197]$, two-tailed). **l**, Sensory-association mean rank axis (Pearson's $r(248) = -0.606$, $p_{spin} = 0.004$, CI = $[-0.679, -0.521]$, two-tailed). **m**, PET tracer binding (BPnd) to MOR (mu-opioid receptor; Pearson's $r(248) = -0.515$, $p_{spin} = 0.025$, CI = $[-0.601, -0.417]$, two-tailed). Source data are provided as a Source Data file.

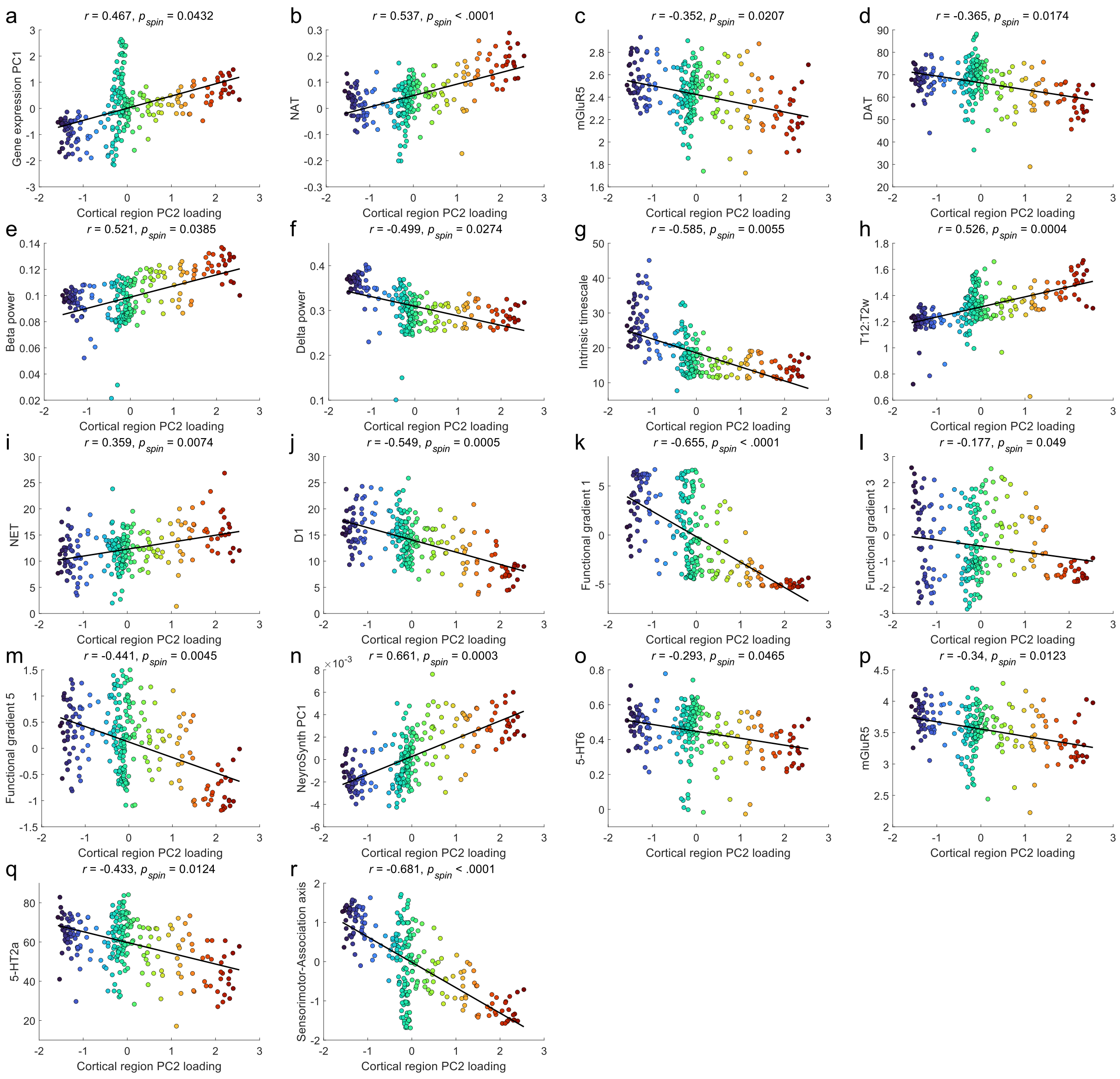


**Fig. S17. Significant relationships between PC2 cortical loadings and cortical measures from the *neuromaps* toolbox.** Points are coloured according to their cortical region PC2 loading. The black line indicates the line of best fit. Significant relationships were detected via a spin-test ($p_{spin} < 0.05$; two-tailed) for the following cortical features: **a**, PC1 of genes in the Allen Human Brain Atlas (Pearson's $r(248) = 0.467$, $p_{spin} = 0.0432$, CI = $[0.364, 0.559]$, two-tailed). **b**, PET tracer binding (BPnd) to NET (norepinephrine transporter; Pearson's $r(248) = 0.537$, $p_{spin} < 0.0001$, CI = $[0.442, 0.62]$, two-tailed). **c**, PET tracer binding (BPnd) to mGluR5 (glutamate receptor; Pearson's $r(248) = -0.352$, $p_{spin} = 0.0207$, CI = $[-0.456, -0.238]$, two-tailed). **d**, PET tracer binding (BPnd) to GABAa (GABA receptor; Pearson's $r(248) = -0.365$, $p_{spin} = 0.0174$, CI = $[-0.468, -0.252]$, two-tailed). **e**, MEG beta (15-29 Hz) power distribution from the Human Connectome Project S1200 release (Pearson's $r(248) = 0.521$, $p_{spin} = 0.0385$, CI = $[0.424, 0.606]$, two-tailed). **f**, MEG delta (2-4 Hz) power distribution from the Human Connectome Project S1200 release (Pearson's $r(248) = -0.499$, $p_{spin} = 0.0274$, CI = $[-0.587, -0.40]$, two-tailed). **g**, MEG intrinsic timescale from the Human Connectome Project S1200 release (Pearson's $r(248) = -0.585$, $p_{spin} = 0.0055$, CI = [-0.661, -0.497], two-tailed). **h**, MRI T1w/T2w ratio from the Human Connectome Project S1200 release (Pearson's $r(248) = 0.526$, $p_{spin} = 0.0004$, CI = $[0.429, 0.61]$, two-tailed). **i**, PET tracer binding (BPnd) to NET (norepinephrine transporter; Pearson's $r(248) = 0.359$, $p_{spin} = 0.0074$, CI = $[0.245, 0.462]$, two-tailed). **j**, PET tracer binding (BPnd)

to D1 (dopamine receptor; Pearson's $r(248) = -0.549$, $p_{spin} = 0.0005$, CI = $[-0.63, -0.456]$, two-tailed). **k**, Diffusion map embedding gradient 1 of group-averaged functional connectivity (Pearson's $r(248) = -0.655$, $p_{spin} < 0.0001$, CI = $[-0.721, -0.578]$, two-tailed). **l**, Diffusion map embedding gradient 3 of group-averaged functional connectivity (Pearson's $r(248) = -0.177$, $p_{spin} = 0.049$, CI = $[-0.295, -0.0546]$, two-tailed). **m**, Diffusion map embedding gradient 5 of group-averaged functional connectivity (Pearson's $r(248) = -0.441$, $p_{spin} = 0.0045$, CI = $[-0.536, -0.336]$, two-tailed). **n**, PC1 of Neurosynth terms in the Cognitive Atlas (123 terms total; Pearson's $r(248) = 0.661$, $p_{spin} = 0.0003$, CI = $[0.585, 0.726]$, two-tailed). **o**, PET tracer binding (BPnd) to 5-HT6 (serotonin receptor; Pearson's $r(248) = -0.293$, $p_{spin} = 0.0465$, CI = $[-0.403, -0.176]$, two-tailed). **p**, PET tracer binding (BPnd) to mGluR5 (glutamate receptor; Pearson's $r(248) = -0.34$, $p_{spin} = 0.0123$, CI = $[-0.446, -0.226]$, two-tailed). **q**, PET tracer binding (BPnd) to 5-HT2a (serotonin receptor; Pearson's $r(248) = -0.433$, $p_{spin} = 0.0124$, CI = $[-0.529, -0.327]$, two-tailed). **r**, Sensory-association mean rank axis (Pearson's $r(248) = -0.681$, $p_{spin} < 0.0001$, CI = $[-0.743, -0.609]$, two-tailed). Source data are provided as a Source Data file.

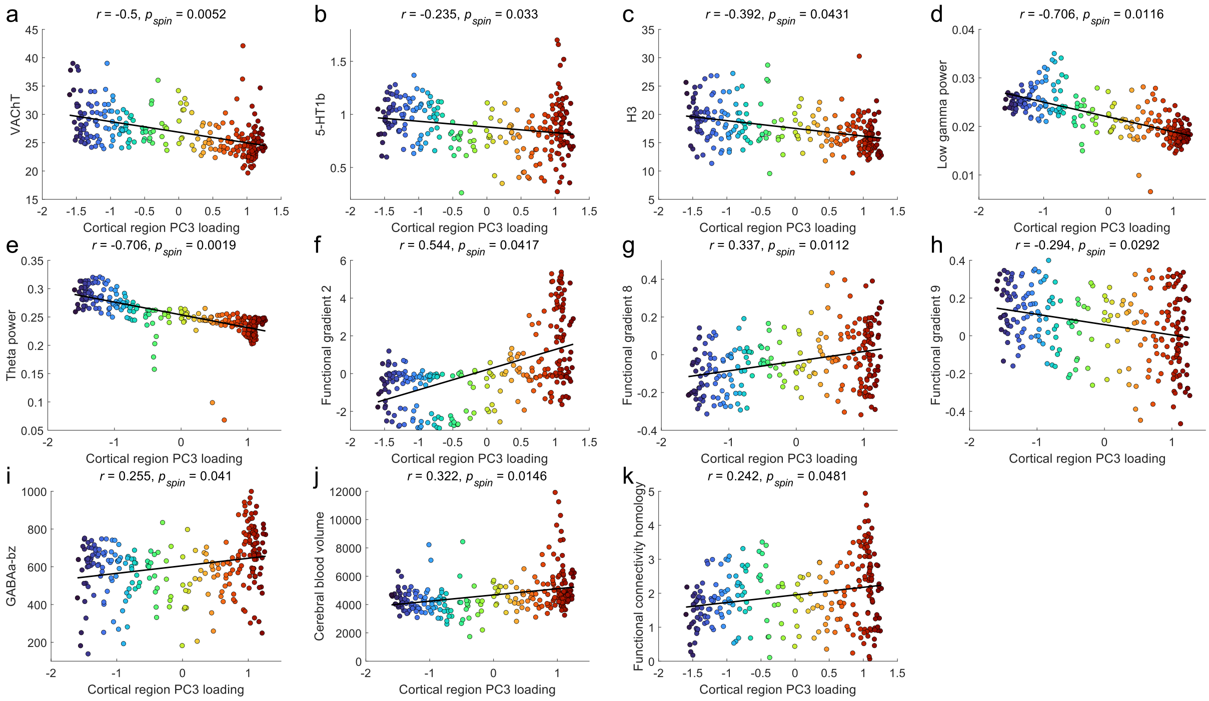


**Fig. S18. Significant relationships between PC3 cortical loadings and cortical measures from the *neuromaps* toolbox.** Points are coloured according to their cortical region PC2 loading. The black line indicates the line of best fit. Significant relationships were detected via a spin-test ($p_{spin} < 0.05$; two-tailed) for the following cortical features: **a**, PET tracer binding (SUVR) to VAChT (acetylcholine transporter; Pearson's $r(248) = -0.5$, $p_{spin} = 0.0052$, CI = $[-0.588, -0.401]$, two-tailed). **b**, PET tracer binding (BPnd) to 5-HT1b (serotonin receptor; Pearson's $r(248) = -0.235$, $p_{spin} = 0.033$, CI = $[-0.349, -0.115]$, two-tailed). **c**, PET tracer binding (Vt) to H3 (histamine receptor; Pearson's $r(248) = -0.392$, $p_{spin} = 0.0431$, CI = $[-0.492, -0.282]$, two-tailed). **d**, MEG low gamma (30-59 Hz) power distribution from the Human Connectome Project S1200 release (Pearson's $r(248) = -0.706$, $p_{spin} = 0.0116$, CI = $[-0.763, -0.638]$, two-tailed). **e**, MEG theta (5-7 Hz) power distribution from the Human Connectome Project S1200 release (Pearson's $r(248) = -0.706$, $p_{spin} = 0.0019$, CI = $[-0.763, -0.638]$, two-tailed). **f**, Diffusion map embedding gradient 2 of group-averaged functional connectivity (Pearson's $r(248) = 0.544$, $p_{spin} = 0.0417$, CI = $[0.45, 0.626]$, two-tailed). **g**, Diffusion map embedding gradient 8 of group-averaged functional connectivity (Pearson's $r(248) = 0.337$, $p_{spin} = 0.0112$, CI = $[0.222, 0.443]$, two-tailed). **h**, Diffusion map embedding gradient 9 of group-averaged functional connectivity (Pearson's $r(248) = -0.294$, $p_{spin} = 0.0292$, CI = $[-0.403, -0.176]$, two-tailed). **i**, PET and autoradiography informed GABAa benzodiazepine binding-site density (GABA receptor; Pearson's $r(248) = 0.255$, $p_{spin} = 0.041$, CI = $[0.135, 0.368]$, two-tailed). **j**, Cerebral blood volume (Pearson's $r(248) = 0.322$, $p_{spin} = 0.0146$, CI = $[0.206, 0.429]$, two-tailed). **k**, Cross-species functional homology (Pearson's $r(248) = 0.242$, $p_{spin} = 0.0481$, CI = $[0.121, 0.355]$, two-tailed). Source data are provided as a Source Data file.

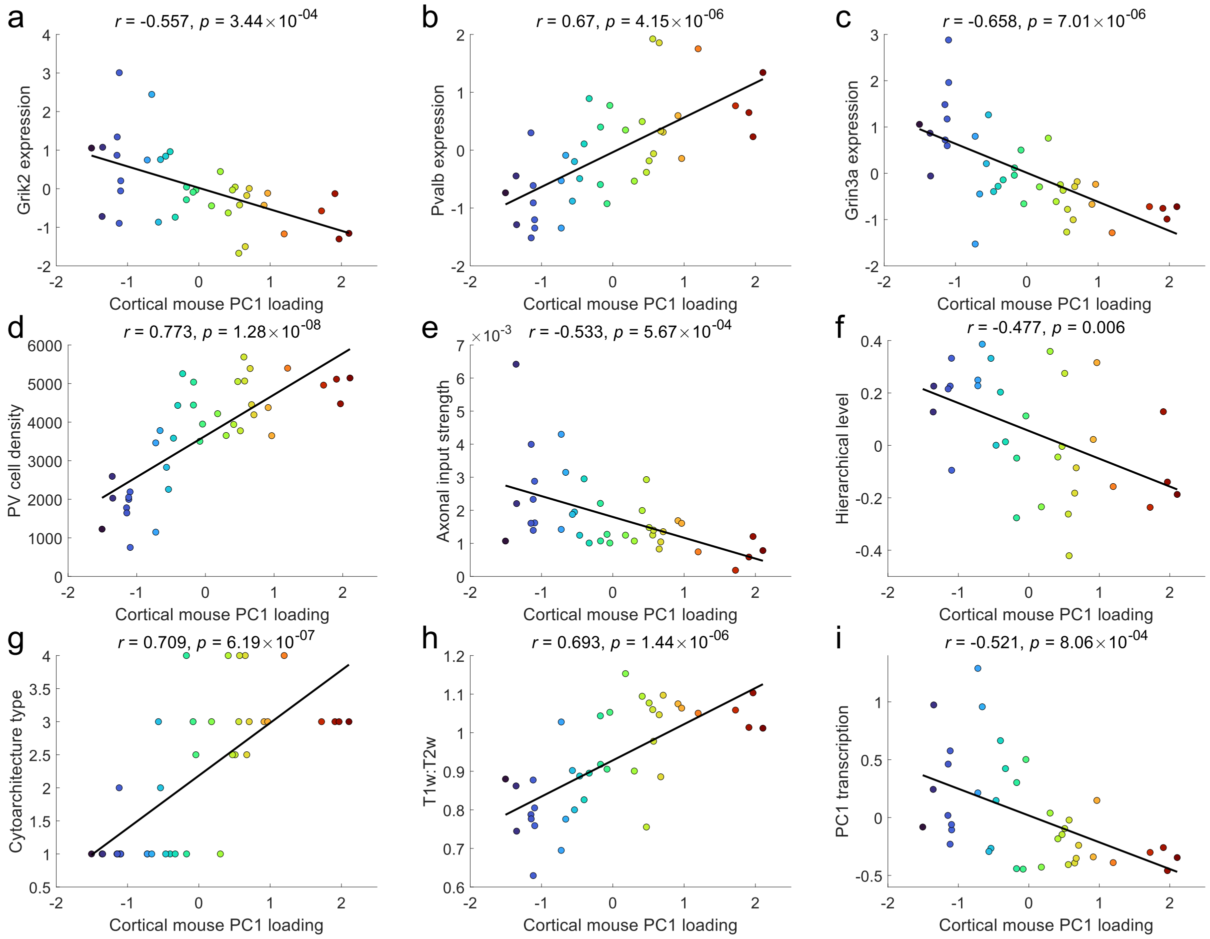


**Fig. S19. Relationship between cortical mouse PC1 loading and measures of mouse cortical hierarchy.** Points are coloured according to their cortical mouse PC1 (mPC1) loading. The black line indicates the line of best fit. These nine measures are considered representative of different cortical features that are illustrative of a hierarchical gradient in the mouse. **a**, Expression of Grik2 (Pearson's $r(35) = -0.557$, $p = 3.44 \times 10^{-04}$, CI = $[-0.746, -0.284]$, two-tailed). **b**, Expression of Pvalb (Pearson's $r(36) = 0.670$, $p = 4.15 \times 10^{-06}$, CI = $[0.446, 0.815]$, two-tailed). **c**, Expression of Grin3a (Pearson's $r(36) = -0.658$, $p = 7.01 \times 10^{-06}$, CI = $[-0.808, -0.429]$, two-tailed). **d**, Mean cell density for parvalbumin-containing (PV) cells (Pearson's $r(36) = 0.773$, $p = 1.28 \times 10^{-08}$, CI = $[0.602, 0.876]$, two-tailed). **e**, Input strength (weighted in-degree) of normalized axonal projection density (Pearson's $r(36) = -0.533$, $p = 5.67 \times 10^{-04}$, CI = $[-0.729, -0.257]$, two-tailed). **f**, Inferred hierarchy from feedforward–feedback laminar projection patterns between cortical and thalamic regions (Pearson's $r(30) = -0.477$, $p = 0.006$, CI = $[-0.708, -0.154]$, two-tailed). **g**, Cytoarchitectonic classification based on regional eulamination (Pearson's $r(36) = 0.709$, $p = 6.19 \times 10^{-07}$, CI = $[0.504, 0.839]$, two-tailed). **h**, Ratio of T1-weighted to T2-weighted (T1w:T2w) images (Pearson's $r(36) = 0.693$, $p = 1.44 \times 10^{-06}$, CI = $[0.479, 0.829]$, two-tailed). **i**, First principal component of mouse cortical gene expression (Pearson's $r(36) = -0.521$, $p = 8.06 \times 10^{-04}$, CI = $[-0.720, -0.241]$, two-tailed). Source data are provided as a Source Data file.

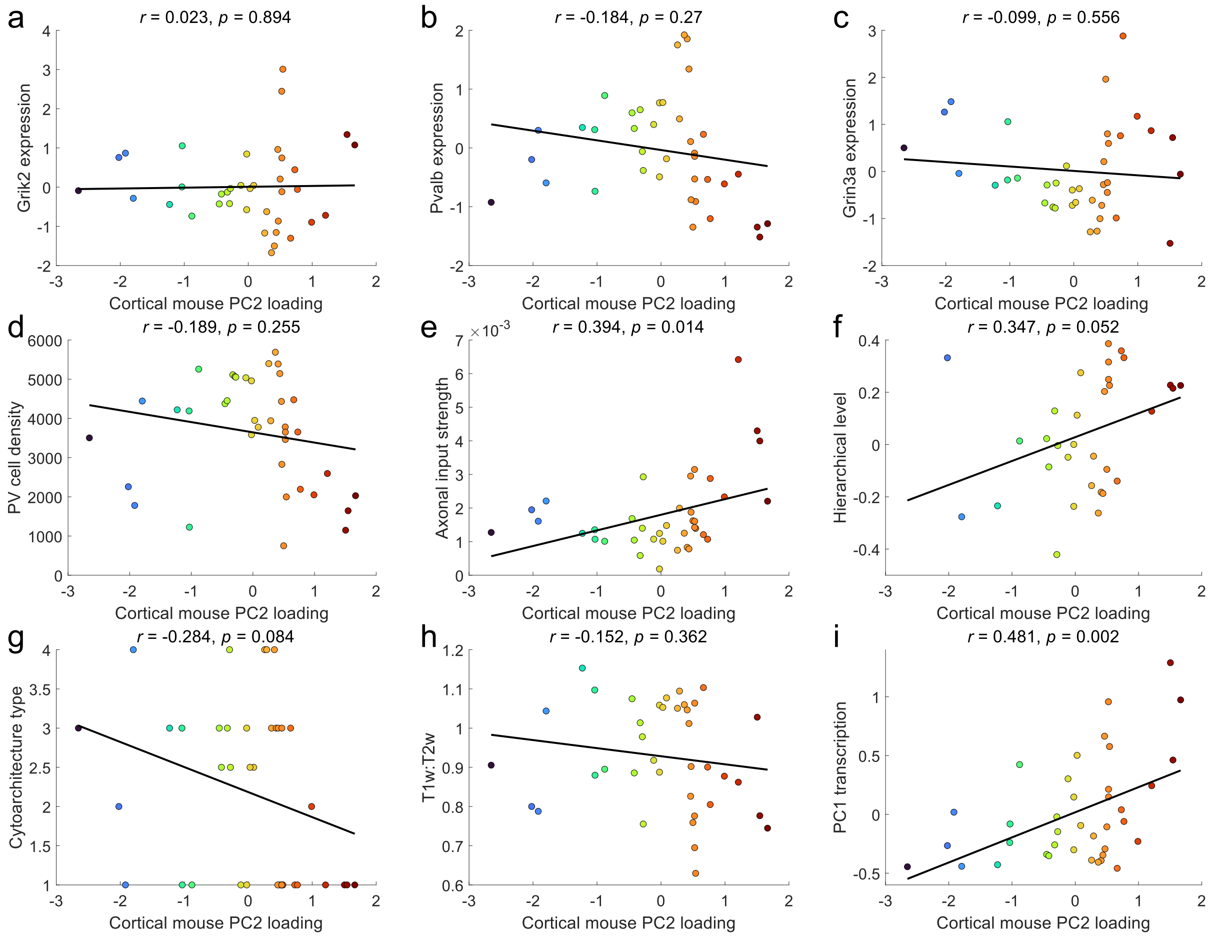


**Fig. S20. Relationship between cortical mouse PC2 loading and measures of mouse cortical hierarchy.** Points are coloured according to their cortical mouse PC2 (mPC2) loading. The black line indicates the line of best fit. These nine measures are considered representative of different cortical features that are illustrative of a hierarchical gradient in the mouse. **a**, Expression of Grik2 (Pearson's $r(35) = 0.023$, $p = 0.894$, CI = $[-0.307, 0.344]$, two-tailed). **b**, Expression of Pvalb (Pearson's $r(36) = -0.184$, $p = 0.70$, CI = $[-0.476, 0.145]$, two-tailed). **c**, Expression of Grin3a (Pearson's $r(36) = -0.099$, $p = 0.556$, CI = $[-0.405, 0.228]$, two-tailed). **d**, Mean cell density for parvalbumin-containing (PV) cells (Pearson's $r(36) = -0.189$, $p = 0.255$, CI = $[-0.480, 0.139]$, two-tailed). **e**, Input strength (weighted in-degree) of normalized axonal projection density (Pearson's $r(36) = 0.394$, $p = 0.0145$, CI = $[0.085, 0.634]$, two-tailed). **f**, Inferred hierarchy from feedforward–feedback laminar projection patterns between cortical and thalamic regions (Pearson's $r(30) = 0.347$, $p = 0.0518$, CI = $[-0.002, 0.621]$, two-tailed). **g**, Cytoarchitectonic classification based on regional eulamination (Pearson's $r(36) = -0.284$, $p = 0.0836$, CI = $[-0.554, 0.039]$, two-tailed). **h**, Ratio of T1-weighted to T2-weighted (T1w:T2w) images (Pearson's $r(36) = -0.152$, $p = 0.362$, CI = $[-0.450, 0.176]$, two-tailed). **i**, First principal component of mouse cortical gene expression (Pearson's $r(36) = 0.481$, $p = 0.0023$, CI = $[0.190, 0.694]$, two-tailed). Source data are provided as a Source Data file.

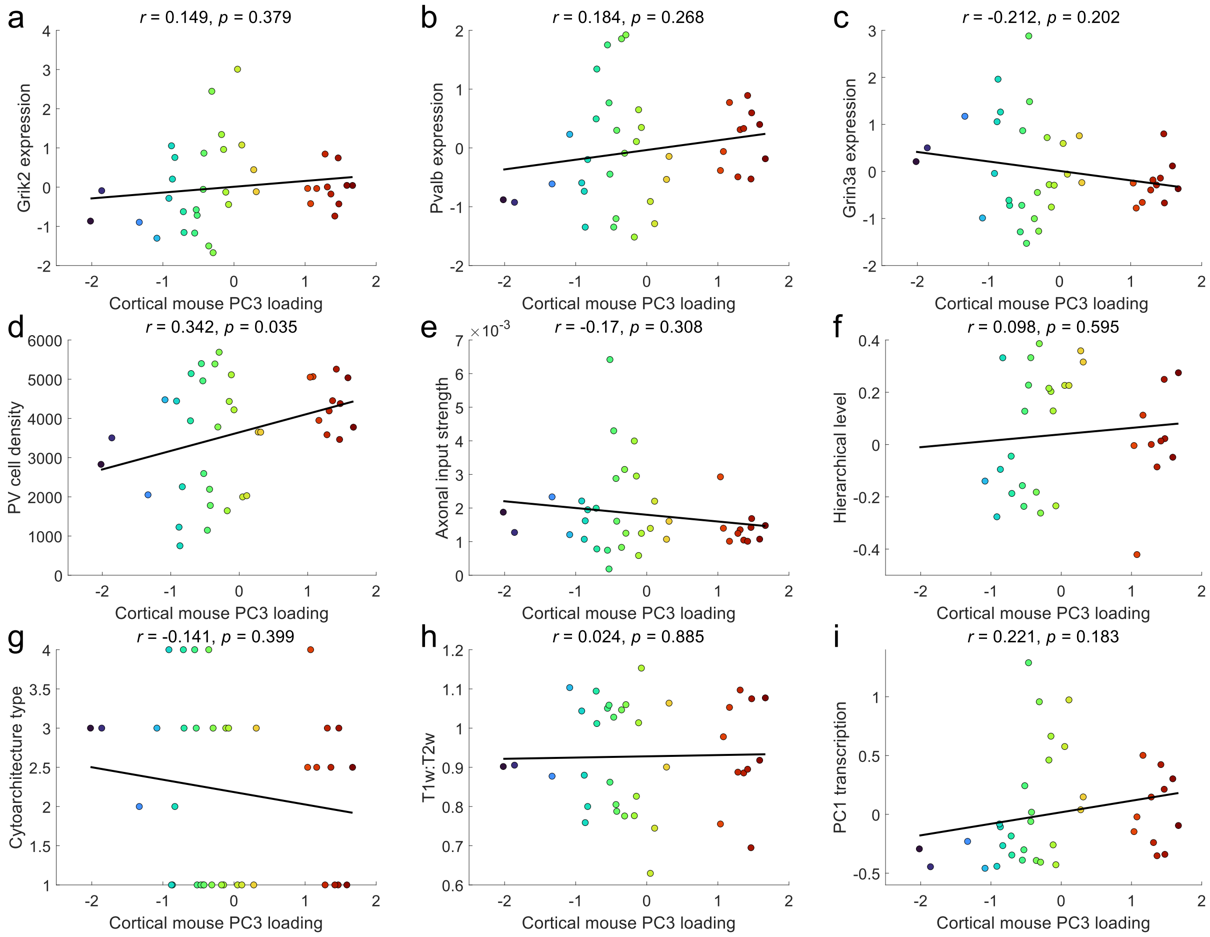


**Fig. S21. Relationship between cortical mouse PC3 loading and measures of mouse cortical hierarchy.** Points are coloured according to their cortical mouse PC3 (mPC3) loading. The black line indicates the line of best fit. These nine measures are considered representative of different cortical features that are illustrative of a hierarchical gradient in the mouse. **a**, Expression of Grik2 (Pearson's $r(35) = 0.149$, $p = 0.379$, CI $= [-0.184, 0.451]$, two-tailed). **b**, Expression of Pvalb (Pearson's $r(36) = 0.184$, $p = 0.268$, CI $= [-0.144, 0.476]$, two-tailed). **c**, Expression of Grin3a (Pearson's $r(36) = -0.212, p = 0.202$, CI $= [-0.498, 0.116]$, two-tailed). **d**, Mean cell density for parvalbumin-containing (PV) cells (Pearson's $r(36) = 0.342$, $p = 0.035$, CI $= [0.025, 0.597]$, two-tailed). **e**, Input strength (weighted in-degree) of normalized axonal projection density (Pearson's $r(36) = -0.170$, $p = 0.308$, CI $= [-0.464, 0.159]$, two-tailed). **f**, Inferred hierarchy from feedforward–feedback laminar projection patterns between cortical and thalamic regions (Pearson's $r(30) = 0.098$, $p = 0.595$, CI $= [-0.260, 0.432]$, two-tailed). **g**, Cytoarchitectonic classification based on regional eulamination (Pearson's $r(36) = -0.141$, $p = 0.399$, CI $= [-0.441, 0.188]$, two-tailed). **h**, Ratio of T1-weighted to T2-weighted (T1w:T2w) images (Pearson's $r(36) = 0.024$, $p = 0.885$, CI $= [-0.298, 0.341]$, two-tailed). **i**, First principal component of mouse cortical gene expression (Pearson's $r(36) = 0.221$, $p = 0.183$, CI $= [-0.107, 0.505]$, two-tailed). Source data are provided as a Source Data file.

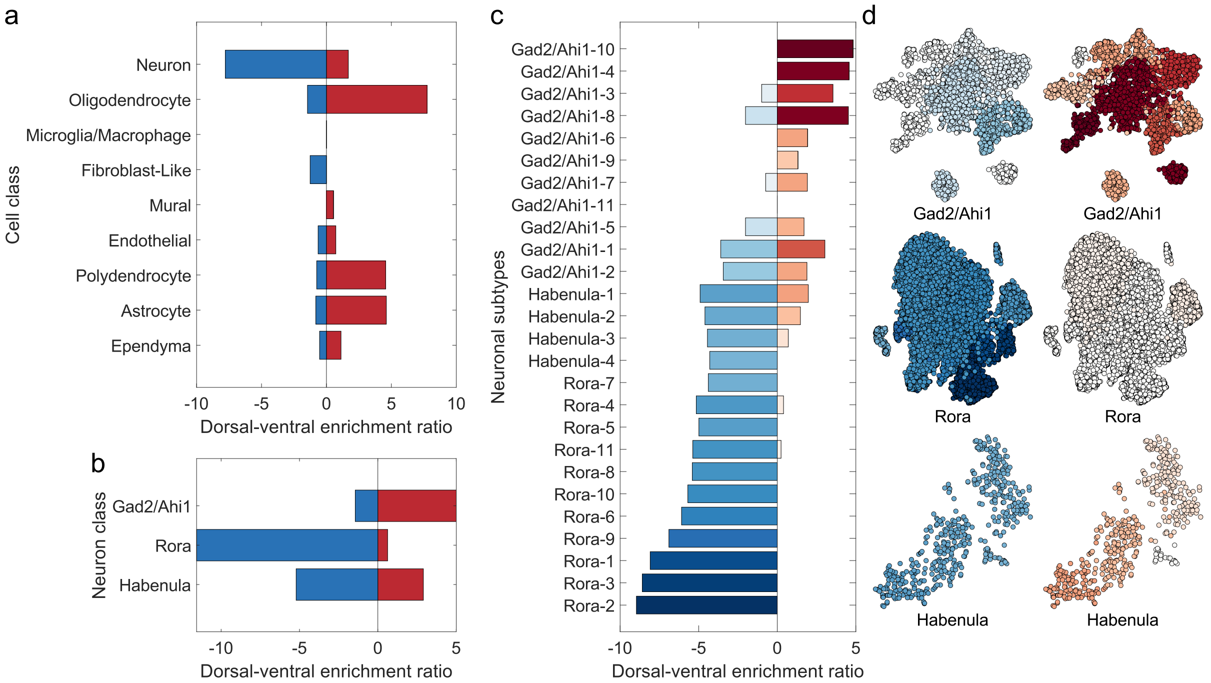


**Figure S22. PC2 cell enrichment results.** **a**, Enrichment ratio for dorsal- and ventral-genes expressed by cell class. The dorsal-ventral enrichment ratio is positive when that cell class is enriched for ventral-genes and negative when enriched for dorsal-genes. **b**, Enrichment ratio for dorsal- and ventral-genes expressed by each neuron class. **c**, Enrichment ratio for dorsal- and ventral-genes expressed by each neuronal subcluster. Subclusters were defined by clustering cells within each neuron class by their pattern of gene expression. In the bar plot, neuronal subclusters are ordered by their summed dorsal-ventral enrichment ratio. **d**, Enrichment ratios projected onto t-SNE plots for the different neuron type subclusters. Cells belonging to each subcluster are coloured according to their enrichment value as indicated in **c** for dorsal- (left) and ventral-genes (right). Source data are provided as a Source Data file.

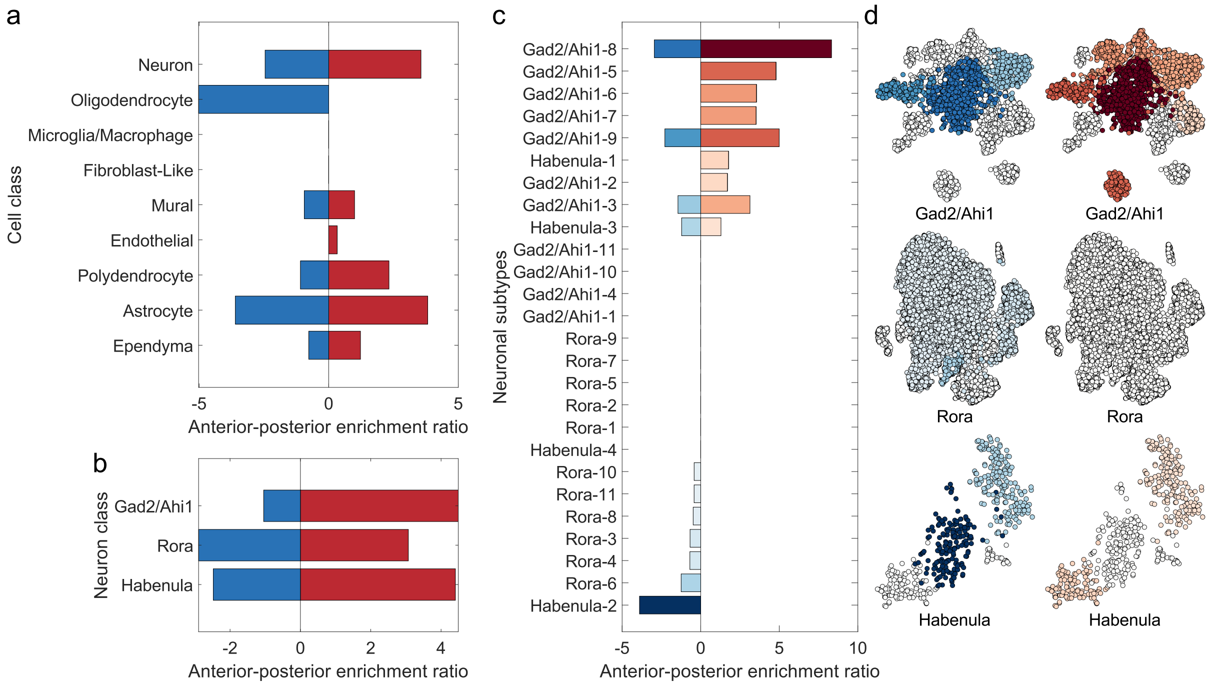


**Figure S23. PC3 cell enrichment results. a**, Enrichment ratio for anterior- and posterior-genes expressed by cell class. The anterior-posterior enrichment ratio is positive when that cell class is enriched for posterior-genes and negative when enriched for anterior-genes. **b**, Enrichment ratio for anterior- and posterior-genes expressed by each neuron class. **c**, Enrichment ratio for anterior- and posterior-genes expressed by each neuronal subcluster. Subclusters were defined by clustering cells within each neuron class by their pattern of gene expression. In the bar plot, neuronal subclusters are ordered by their summed anterior-posterior enrichment ratio. **d**, Enrichment ratios projected onto t-SNE plots for the different neuron type subclusters. Cells belonging to each subcluster are coloured according to their enrichment value as indicated in **c** for anterior- (left) and posterior-genes (right). Source data are provided as a Source Data file.

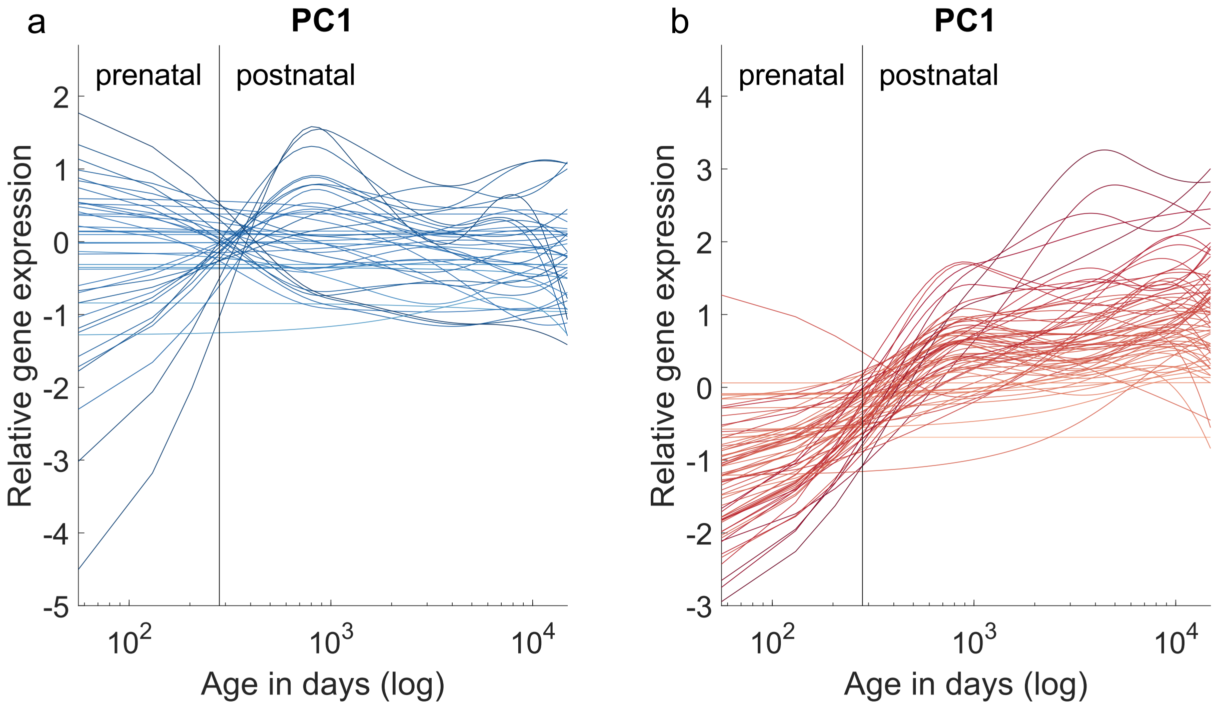


**Fig. S24. Trajectories of gene expression for PC1 enriched genes. a**, Relative gene expression of medial-genes over time. **b**, Relative gene expression of lateral-genes over time. Source data are provided as a Source Data file.

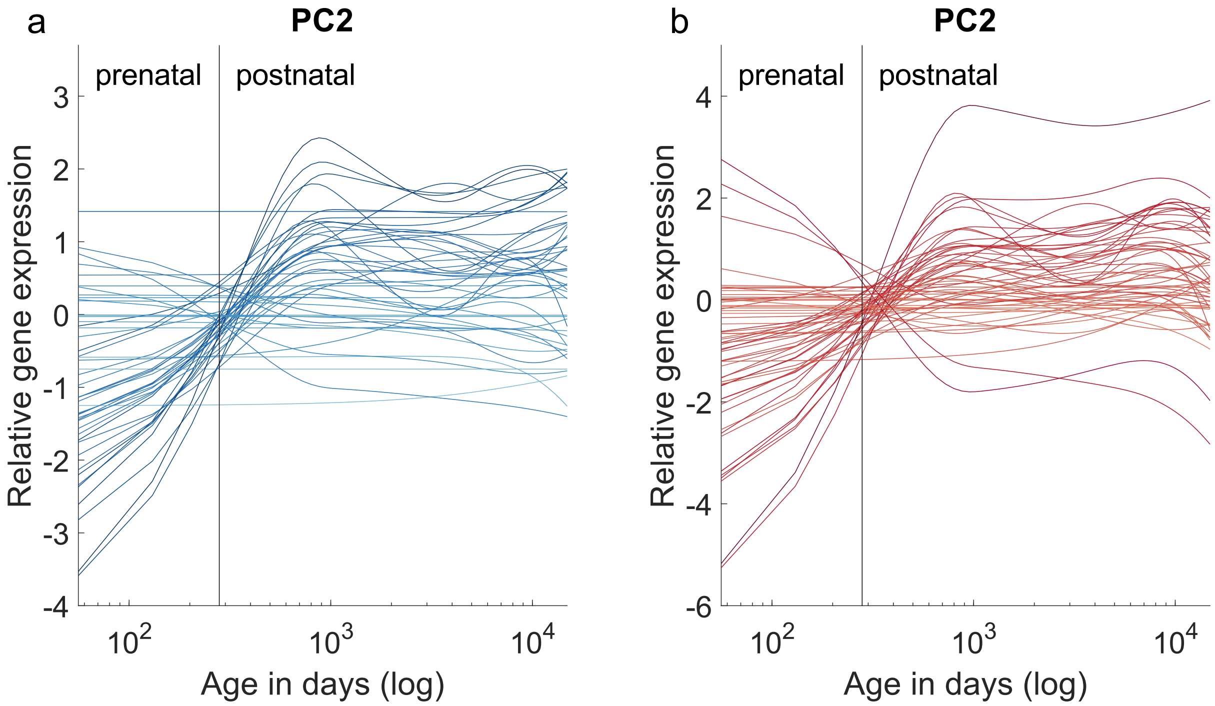


**Fig. S25. Trajectories of gene expression for PC2 enriched genes. a**, Relative gene expression of genes negatively correlated with PC2 over time. **b**, Relative gene expression of genes positively correlated with PC2 over time. Source data are provided as a Source Data file.

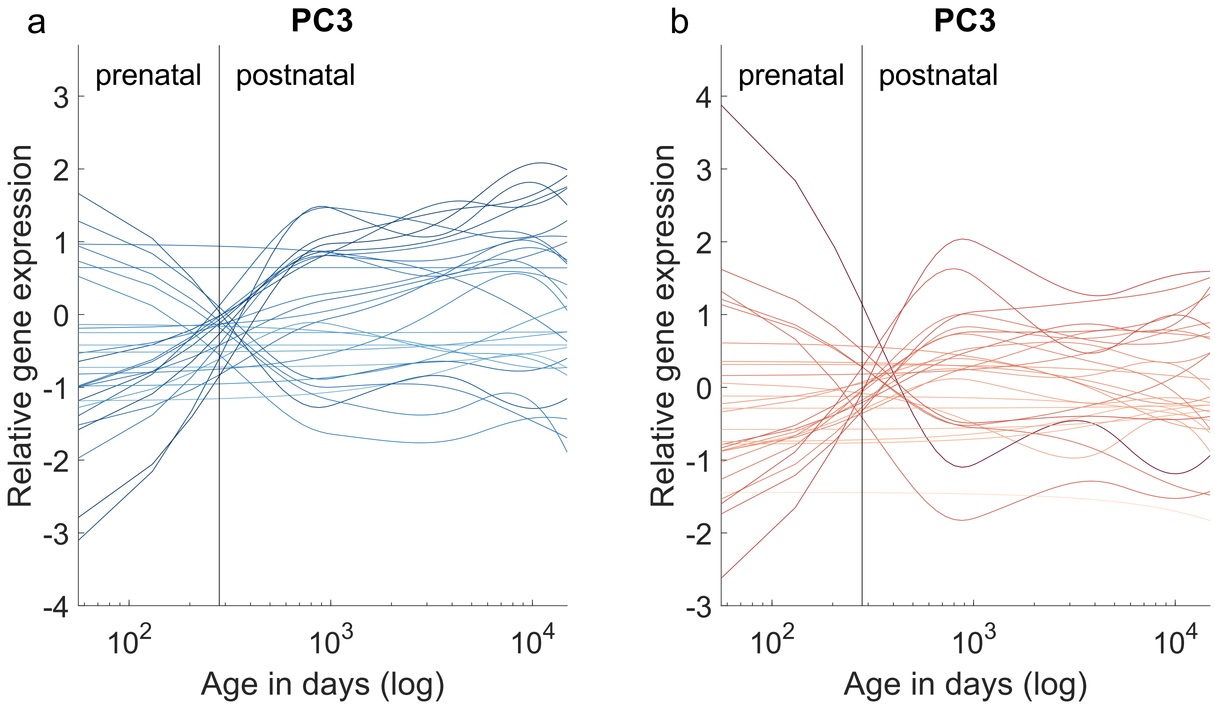


**Fig. S26. Trajectories of gene expression for PC3 enriched genes. a**, Relative gene expression of genes negatively correlated with PC3 over time. **b**, Relative gene expression of genes positively correlated with PC3 over time. Source data are provided as a Source Data file.
